# One high-order ion-coupling mechanism generates multiple SLC1A stoichiometries

**DOI:** 10.64898/2026.11.07.764996

**Authors:** Qianyi Wu, Yan Jiang, Chelsea Briot, Ashkan Fakharzadeh, Morgan M. Dooley, Shashank Pant, Hariprasad Venugopal, Krishna D. Reddy, Emad Tajkhorshid, Renae M. Ryan

## Abstract

Human excitatory amino acid transporter 2 (hEAAT2) is a glial transporter that clears glutamate and aspartate from the extracellular space to terminate neurotransmission. When this essential role is disrupted, excess extracellular glutamate causes excitotoxicity and neurological disease. hEAAT2 transports amino acids into cells against their concentration gradients by coupling uptake to the co-transport of Na^+^ and H^+^, and counter-transport of K^+^. Substrate transport also activates a thermodynamically uncoupled Cl^-^ conductance, whose significance is unknown. Here, to dissect the molecular mechanisms underlying coupled and uncoupled ion transport by EAATs, we leverage the tractable *Drosophila melanogaster* homologue dEAAT2, which transports taurine as well as glutamate and aspartate. By functionally characterising dEAAT2, we confirmed it has features of a canonical EAAT transport cycle, including an uncoupled Cl^-^ conductance. However, it functions an ion-independent electroneutral exchanger, in stark contrast to its ion-coupled electrogenic human counterpart. Cryo-electron microscopy (cryo-EM) and computational simulations confirmed the inability of dEAAT2 to transport cations, but just two point mutations conferred Na^+^, H^+^, and K^+^ dependence, reconstituting the complex ion coupling profile of human transporters in the *Drosophila* exchanger. We also showed that dEAAT2 transports taurine in a complex with Cl^-^, bestowing a unique Cl^-^-dependence on taurine transport, independent of uncoupled Cl^-^ flux. Our findings show that ancestral exchangers in the EAAT family preserve the machinery to achieve complex ion-coupled transport, revealing the molecular basis of high-order ion coupling in hEAATs.

## Main

In the central nervous system, transmitters released into synapses must be recycled back to presynaptic terminals for repeated rounds of neurotransmission. Uptake of transmitters from the extracellular environment is carried out by membrane-bound transporters, including the <u>e</u>xcitatory <u>a</u>mino <u>a</u>cid<u>-t</u>ransporting EAATs, which recycle the majority of glutamate in the human brain and whose dysfunction leads to episodic ataxia^1–4^, glioma^5^, amyotrophic lateral sclerosis^6^, cerebral ischemia^7^, Alzheimer’s disease^8^, and poor recovery from stroke^9^. EAATs are secondary active transporters that belong to the solute carrier 1A (SLC1A) family and utilise energy from the electrochemical gradients of ions to concentrate glutamate and aspartate within cells. Transport of one substrate molecule is tightly coupled to the cotransport of three Na^+^, one H^+^, and the counter-transport of one K^+^, creating an electrogenic transfer of positive charge into the cell^10–12^. In addition, Na^+^ and substrate binding activates a thermodynamically uncoupled Cl^-^ conductance through a pathway that is distinct from the substrate translocation path^13–19^ (Fig.1a).

SLC1A family members have been utilised throughout evolution, including by *Drosophila melanogaster*, which expresses two EAATs in its central nervous system, dEAAT1 and dEAAT2^20^. Although dEAAT1 transports glutamate and aspartate with similar affinity^21^, dEAAT2 favours aspartate over glutamate and transports taurine with high-affinity^22^, despite its 40% sequence identity with hEAAT2^20^. Taurine is the second most abundant amino acid in both insect^23^ and mammalian brains^24^. In mammals, taurine acts as an inhibitory neurotransmitter that regulates cell volume^25,26^, contributes to nervous system development^27^, and protects against excitotoxicity in the hippocampus^28^. Taurine is transported by members of the solute carrier 6 (SLC6) family in mammals, however there are no homologues of this family in insects, which instead recover taurine via dEAAT2^20,22^. Indeed, recent work has demonstrated an important role for dEAAT2 in hearing^29^, sleep modulation, and metabolism^30^.

Extensive functional and structural studies of mammalian and prokaryotic SLC1A transporters have shown that they operate with an elevator-like mechanism, in which a transport domain moves across the membrane with respect to a scaffold domain^31–33^. However, the molecular mechanisms responsible for coupled and uncoupled ion transport remain enigmatic. Thus, although three binding sites for Na^+^ (Na1-3) have been identified^34–36^, the H^+^- and K^+^-binding sites, whose locations are crucial for understanding coupled transport, remain controversial^37–40^. Like Na^+^, H^+^ favours the forward transport (uptake) of substrates, however K^+^ binding reorientates the transporter from its inward-facing state (IFS) back to its outward-facing state (OFS) in preparation for another transport cycle^41^. Under pathological conditions when Na^+^ and K^+^ gradients are disrupted, such as following stroke, reverse transport can lead to reduced extracellular glutamate clearance or even glutamate efflux from cells^42^. Interestingly, a highly conserved glutamate residue in transmembrane helix 7 (TM7) of hEAATs is replaced by an uncharged glutamine in family members that are not H^+^-dependent (Extended Data Fig. 1), suggesting this may be the site of H^+^ binding^33^. In addition, a double alanine motif in the tip of hairpin 2 (HP2), which is thought to be involved in K^+^ binding^43^, is conserved in K^+^-dependent EAATs but absent in the K^+^-independent prokaryotic homolog Glt_Ph_ (Extended Data Fig. 1). Because dEAAT2 possesses an uncharged isoleucine residue in the conserved region of TM7 and a serine residue in place of the first alanine in the HP2 motif, we predicted that this insect homologue could be used to explore cation coupling in SLC1A transporters.

In this study, we characterised the full transport mechanism of dEAAT2, revealing that it is a Na^+^/H^+^/K^+^-independent aspartate and taurine electroneutral exchanger with an uncoupled Cl^-^ conductance. Interestingly, we found that substrate transport is positively modulated by Na^+^, even though cations aren’t required for transport activity, and that taurine transport has a strict requirement for Cl^-^, independent of the uncoupled Cl^-^ conductance. Using a combination of functional assays, cryo-EM and computational simulations, we showed that just two mutations in dEAAT2 could reconstitute canonical, human-like ion coupling, revealing the molecular basis of Na^+^, H^+^, and K^+^ coupling in the SLC1A family. Our data showcase the utility of dEAAT2 as a tool to investigate transport mechanisms in vertebrate SLC1A family members and explain how some function as transporters and others as exchangers.

### dEAAT2 is an electroneutral exchanger

To characterise the stoichiometry of dEAAT2, we expressed the transporter in *Xenopus laevis* oocytes and compared its functional properties to those of hEAAT2. When oocytes expressing hEAAT2 were incubated with radiolabelled substrates, we observed robust transport of _L_-[^3^H]glutamate (Fig. 1b) but no [^3^H]taurine uptake, as expected (Fig. 1c). In contrast, dEAAT2 was a relatively poor transporter of glutamate (Fig.1b) but facilitated robust uptake of [^3^H]taurine (Fig. 1c). Both hEAAT2 and dEAAT2 transported _L_-[^3^H]aspartate, and addition of excess unlabelled _L_-glutamate suppressed this uptake by 62-64%, suggesting that _L_-glutamate and _L_-aspartate compete for the same binding site in both transporters (Fig. 1d). Unlabelled taurine did not inhibit _L_-[^3^H]aspartate uptake by hEAAT2 but suppressed its uptake by dEAAT2 to almost background levels (Fig. 1d), underscoring the different substrate selectivity of human and insect transporters. As expected, the aspartate-based SLC1A inhibitor _DL_-TBOA^40^ was effective in abolishing substrate uptake and ion transport by both dEAAT2 and hEAAT2 (Supplementary Data Fig. 1).

**Figure 1.**
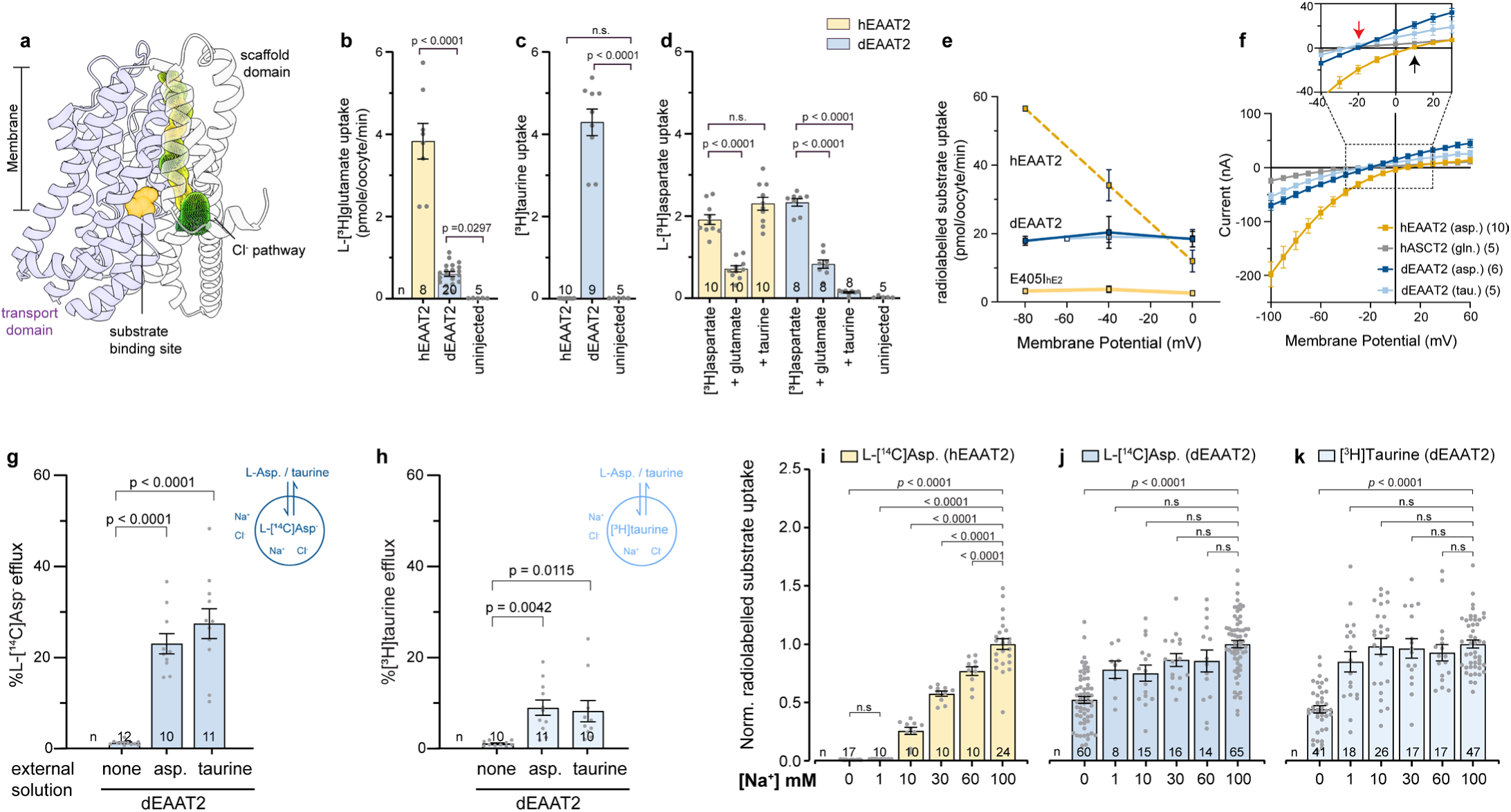
dEAAT2 transports taurine, aspartate, and glutamate. (**a**) The chloride-conducting state (ClCS) of SLC1A archaeal homolog Glt_Ph_ (PDB 6WYK)^15^ showing the substrate binding site (yellow) and Cl^-^ conducting pathway (mesh) at the interface of the transport domain (purple) and scaffold domain (white). Uptake of 10 µM _L_-[^3^H]glutamate (**b**), [^3^H]taurine (**c**), and _L_-[^3^H]aspartate (**d**) by hEAAT2- and dEAAT2-expressing oocytes. Background uptake from control (uninjected) oocytes shown for each radiolabelled substrate. (**d**) _L_-[^3^H]aspartate uptake in the absence and presence of 1 mM unlabelled glutamate or taurine. (**e**) Radiolabelled substrate uptake by hEAAT2, dEAAT2 and hEAAT2 mutation E405I_hE2_ over a range of membrane potentials. hEAAT2 substrate was _L_-[^3^H]aspartate, dEAAT2 substrates were [^3^H]taurine (open symbols) and _L_-[^3^H]aspartate (closed symbols). Data are mean ± SEM (n=3, except dEAAT2 -80mV n=5, E405I_hE2_ n=4). (**f**) *E_rev_* from current-voltage (I-V) relationships in response to application of 1 mM aspartate (hEAAT2 and dEAAT2), 1 mM taurine (dEAAT2), and 1 mM glutamine (hASCT2). Upper panel, magnified view of *E_rev_*. Efflux of preloaded _L_-[^14^C]aspartate (**g**) or [^3^H]taurine (**h**) from dEAAT2-expressing oocytes in the absence and presence of 300 µM unlabelled _L_-aspartate or taurine to initiate exchange. Background efflux measured from control oocytes. _L_-[^14^C]aspartate uptake via hEAAT2 (**i**) and _L_-[^14^C]aspartate (**j**) and [^3^H]taurine (**k**) uptake via dEAAT2 were examined in a range of external Na^+^ concentrations. Responses from individual oocytes shown as grey dots. Data obtained from at least two different batches of oocytes and presented as mean ± SEM for the indicated number of cells (n) unless otherwise stated. Data analysed using ordinary One-Way ANOVA with Dunnett’s multiple comparison tests by GraphPad Prism 10; p values as indicated (n.s., not significant).

We also examined the transport mechanism utilised by dEAAT2 by monitoring radiolabelled substrate uptake over a range of membrane potentials. The electrogenic nature of _L_-aspartate transport via hEAAT2 was observed as a reduction in _L_-[^14^C]aspartate uptake as membrane potential was depolarised (Fig. 1e). In contrast, transport of both _L_-[^14^C]aspartate and [^3^H]taurine by dEAAT2 was similar at membrane potentials between - 80 mV and 0 mV, suggesting that dEAAT2 is an electroneutral transporter. As expected, the overall substrate-elicited conductance of hEAAT2 had a positive reversal potential (*E_rev_*) in our recording conditions (Fig. 1f, black arrow), due to the combination of inward ion-coupled transport and uncoupled Cl^-^ flux that reverses close to the equilibrium potential for Cl^-^ (*E_Cl-_*; -24 mV)^44^. In contrast, the conductance elicited by _L_-aspartate and taurine in dEAAT2 reversed very close to *E_Cl-_*, resembling that generated by glutamine in the electroneutral exchanger hASCT2^45,46^, whose current is carried solely by its uncoupled Cl^-^ conductance (Fig. 1f, red arrow). Upon loading dEAAT2-expressing oocytes with either _L_-[^14^C]aspartate or [^3^H]taurine in the presence of external unlabelled substrate, we observed robust efflux in both cases (Fig. 1g, h) and dEAAT2 can transport both aspartate and taurine in the absence of external Na^+^ (Fig. 1j-k). This contrasts with hEAAT2, which has a strict requirement for external Na^+^ (Fig. 1i). These data demonstrate that dEAAT2 functions as a Na^+^-independent electroneutral aspartate/taurine exchanger with an associated Cl^-^ conductance – the first example of an acidic amino exchanger in the SLC1A family.

### Taurine transport by dEAAT2 is Cl^-^-dependent

In hEAATs, the Cl^-^ conductance has the following anion permeability sequence: F^-^<< Cl^-^< Br^-^< I^-^< NO_3_^-^. Because NO_3_^-^ is about ten times more permeable than Cl^-^, substrate-elicited outward currents in oocytes expressing hEAAT2 were greater when Cl^-^was replaced by NO_3_^-^ in the recording buffer (Fig. 2a). In contrast, dEAAT2 currents were reduced and inwardly directed in the presence of NO_3_^-^, suggesting that either the anion conductance of dEAAT2 has a different permeability sequence or that NO_3_^-^ inhibits taurine transport. To distinguish between these two possibilities, we measured uptake of radiolabelled substrates in the presence of various anions. In hEAAT2, the rate of _L_-[^14^C]aspartate transport was unaffected by the choice of anion, including impermeant anions such as methanesulfonate (MES^-^) (Fig. 2b). In addition, the *K_m_* for _L_-aspartate was similar in the presence of different anions (Extended Data Table 1) due to the uncoupled nature of the hEAAT2 Cl^-^ conductance. In contrast, [^3^H]taurine uptake via dEAAT2 sequentially decreased as Cl^-^ was substituted with Br^-^, I^-^, and NO_3_^-^, and was absent in the presence of MES^-^ or the highly electronegative F^-^ (Fig. 2d). This suggests that [^3^H]taurine transport is dependent on Cl^-^ and that the ability to support [^3^H]taurine transport decreases as the size of the anion increases. This anion dependence was also apparent from *K_m_* measurements for taurine, which became larger (indicating reduced affinity) as anion size increased (Extended Data Fig.2a-c and Extended Data Table 1). Interestingly, the effect of anion size on _L_-[^14^C]aspartate transport by dEAAT2 was less remarkable, the rate being reduced by only 46% and 56% in I^-^ and NO_3_^-^ buffers, respectively (Fig. 2c), and the *K_m_* increasing about twofold in the presence of Br^-^ and I^-^, respectively (Extended Data Table 1). Moreover, when Cl^-^was replaced by MES^-^ or F^-^, _L_-[^14^C]aspartate uptake was unaffected, suggesting that _L_-aspartate transport does not require Cl^-^ but that I^-^ and NO_3_^-^ may affect _L_-[^14^C]aspartate binding. Indeed, _L_-[^14^C]aspartate uptake was unaffected by changes in external Cl^-^. However, [^3^H]taurine transport displayed a clear dependence on Cl^-^ with an apparent affinity of 10.5 mM (Fig. 2e).

**Figure 2.**
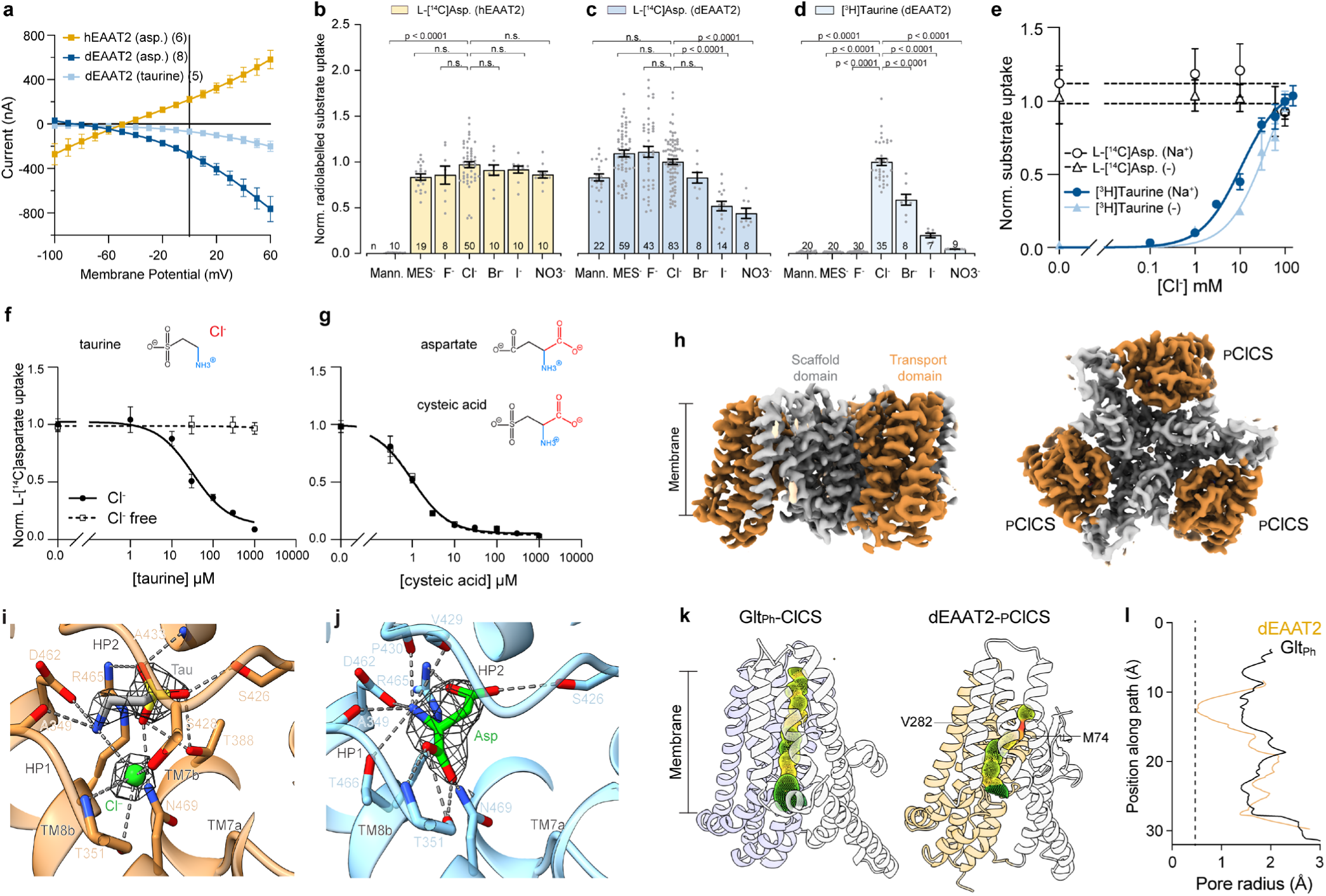
Chloride dependence of hEAAT2 and dEAAT2. (**a**) IV relationships generated by application of aspartate or taurine to hEAAT2- and dEAAT2-expressing oocytes in a NO_3_^-^ buffer. _L_-[^14^C]aspartate uptake by (**b**) hEAAT2 or (**c**) dEAAT2 and (**d**) [^3^H]taurine uptake by dEAAT2 with the indicated permeable anions or without ions in _D_-mannitol buffer (Mann.). Responses from individual oocytes shown as grey dots. Data obtained from at least two different batches of oocytes and presented as mean ± SEM for the indicated number of cells (n). Data analysed using ordinary One-Way ANOVA with Dunnett’s multiple comparison tests by GraphPad Prism 10; p values as indicated (n.s., not significant). (**e**) Cl^-^ dose-response curves in dEAAT2-expressing oocytes using buffers containing 10 µM [^3^H]taurine (blue) or _L_-[^14^C]aspartate (black) in the absence (triangle) and presence (circle) of Na^+^. Apparent affinity (K*_m_*) for Cl^-^ could only be determined for taurine, in the presence of Na^+^ 10.5 mM (7.4 – 15.1) and in the absence of Na^+^ 40.1 mM (22.3 – 80.5). Brackets indicate 95% confidence intervals. _L_-[^14^C]aspartate competition assay with various concentrations of (**f**) taurine or (**g**) cysteic acid in the absence (open symbols) or presence (closed symbols) of Cl^-^, using MES^-^ as a substitute for Cl^-^. See Supplementary Table 2 for exact n. **(h)** A Cryo-EM density map of dEAAT2 bound to taurine at 2.4 Å resolution, shown in side (left) and top-down (right) view. Each protomer is coloured by domain: transport (orange), scaffold (grey). All three protomers adopt the pre-open chloride-conducting state conformation (_P_ClCS). Close-up of the taurine (**i**) or aspartate (**j**) binding pocket in the _P_ClCS conformation. Cryo-EM density for substrates shown in mesh. (**k**) Taurine-bound cryoEM structure shows a dominant _P_ClCS, similar to the ClCS of Glt_Ph_. The conducting pathway is drawn by the Hole plugin in Coo revealing a constriction point for dEAAT2 formed by M74 and V282 (red). (**l**) The pore radius of Glt_Ph_ (ClCS) and dEAAT2 (_P_ClCS).

At physiological pH, taurine exists as a neutral zwitterion, whereas aspartate is negatively charged due to a second carboxyl group on the α-carbon. Assuming taurine and aspartate occupy the same binding site (Fig. 1d), we hypothesised that Cl^-^ may substitute for the second carboxyl group on aspartate, allowing taurine to bind to the same substrate-binding pocket of dEAAT2. To test this, we examined the Cl^-^ dependence of a series of _L_-aspartate and taurine derivatives using an _L_-[^14^C]aspartate competition assay. This revealed that taurine inhibits _L_-[^14^C]aspartate uptake with an apparent affinity of 32.3 µM, but only in the presence of Cl^-^ (Fig. 2f). This requirement for Cl^-^ was observed for all compounds lacking a second carboxyl group (Extended Data Fig. 2d-f and Extended Data Table 2). However, we did not find any Cl^-^ dependence for compounds containing a second negatively charged moiety, including cysteic acid, which inhibits _L_-[^14^C]aspartate uptake in the absence as well as the presence of Cl^-^ (Fig. 2g, Extended Data Fig. 2g-i and Extended Data Table 2).

We validated our hypothesis by solving the structure of dEAAT2 with bound taurine. In the presence of saturating concentrations of Na^+^ and taurine, we observed symmetrical dEAAT2 trimers in a novel intermediate conformation (Fig. 2h), featuring a Cl^-^-conducting pathway with a constriction at a hydrophobic cluster formed by M74 and V282, equivalent to the extracellular gate of the Cl^-^ channel in hEAAT1 and Glt_Ph_^15,47^ (Fig. 2k,l; Extended Data Fig. 3a,b; Supplementary Data Fig. 2-4; Supplementary Table 1). We named this conformation the pre-open Cl^-^-conducting state (_P_ClCS) as it would require an additional ∼7° rotation of the transport domain to open the Cl^-^ channel. Aspartate-bound dEAAT2 monomers were also found in _P_ClCS, as well as an outward-facing closed state (OF_C_S) (Supplementary Data Fig. 5-6; Supplementary Table 1).

Cryo-EM maps revealed two distinct non-protein densities in the taurine-binding pocket, corresponding to a taurine molecule and a distinct spherical density (Fig. 2i), in contrast to the aspartate-binding pocket where a single non-protein density was observed (Fig. 2j). Because our functional data demonstrated that taurine transport is dependent on extracellular Cl^-^ (Fig 2d) and Cl^-^ is the only anion present in the purification and imaging buffers, we modelled this spherical density as a Cl^-^ ion. Taurine is stabilised by extensive hydrogen bonds with surrounding residues. Its sulfonic acid interacts with residues on TM8 (R465, N469), HP2 (S426, A433), and TM7 (T388), while the amino group forms hydrogen bonds with TM8 (D462) and HP1 (A349). The modelled Cl^-^ coordinates the amino group of taurine as well as polar residues on HP1 (T351) and HP2 (S428), mimicking the second carboxylate of aspartate (Fig. 2i,j). Thus, the uncoupled Cl^-^ pathway in the *Drosophila* homologue resembles that in the human transporter, but dEAAT2 co-opts Cl^-^ to transport taurine into cells as a taurine:Cl^-^ complex that occupies the same binding position as aspartate, resulting in a unique transport dependence on extracellular Cl^-^ that is independent of the uncoupled anion conductance (Fig. 1a).

### I392E/S426A_dE2_ confers H^+^/K^+^ dependence

Having established the electroneutrality of substrate exchange by dEAAT2, we sought to determine whether this exchange is driven by ion flux, albeit via a net neutral mechanism. We first examined a conserved TM7 glutamate residue that has been proposed to facilitate H^+^ coupling in hEAATs (E405_hE2_)^41,48,33^. This glutamate residue is replaced by an uncharged glutamine in the H^+^/K^+^-independent transporters ASCT1-2 and Glt_Ph_^46,49^ and an uncharged isoleucine in dEAAT2 (I392_dE2_; Extended Data Fig. 1). To explore the H^+^ dependence of substrate transport, we determined dEAAT2 activity by measuring _L_-[^14^C]aspartate uptake in either the presence of symmetrical H^+^ or an inwardly directed H^+^ gradient. Unlike hEAAT2, which displayed increased _L_-[^14^C]aspartate uptake in the presence of an inward H^+^ gradient, _L_-[^14^C]aspartate uptake by dEAAT2 was similar in all conditions (Fig. 3a). However, mutation of I392 to a negatively charged glutamate (I392E_dE2_) introduced H^+^ dependence into dEAAT2, and conversely, mutation of E405 to an uncharged glutamine (E405Q_hE2_) abolished H^+^ dependence in hEAAT2^48^. These data support a role for E405_hE2_ as the H^+^ carrier in hEAAT2 and explain why dEAAT2 is H^+^-independent.

**Figure 3.**
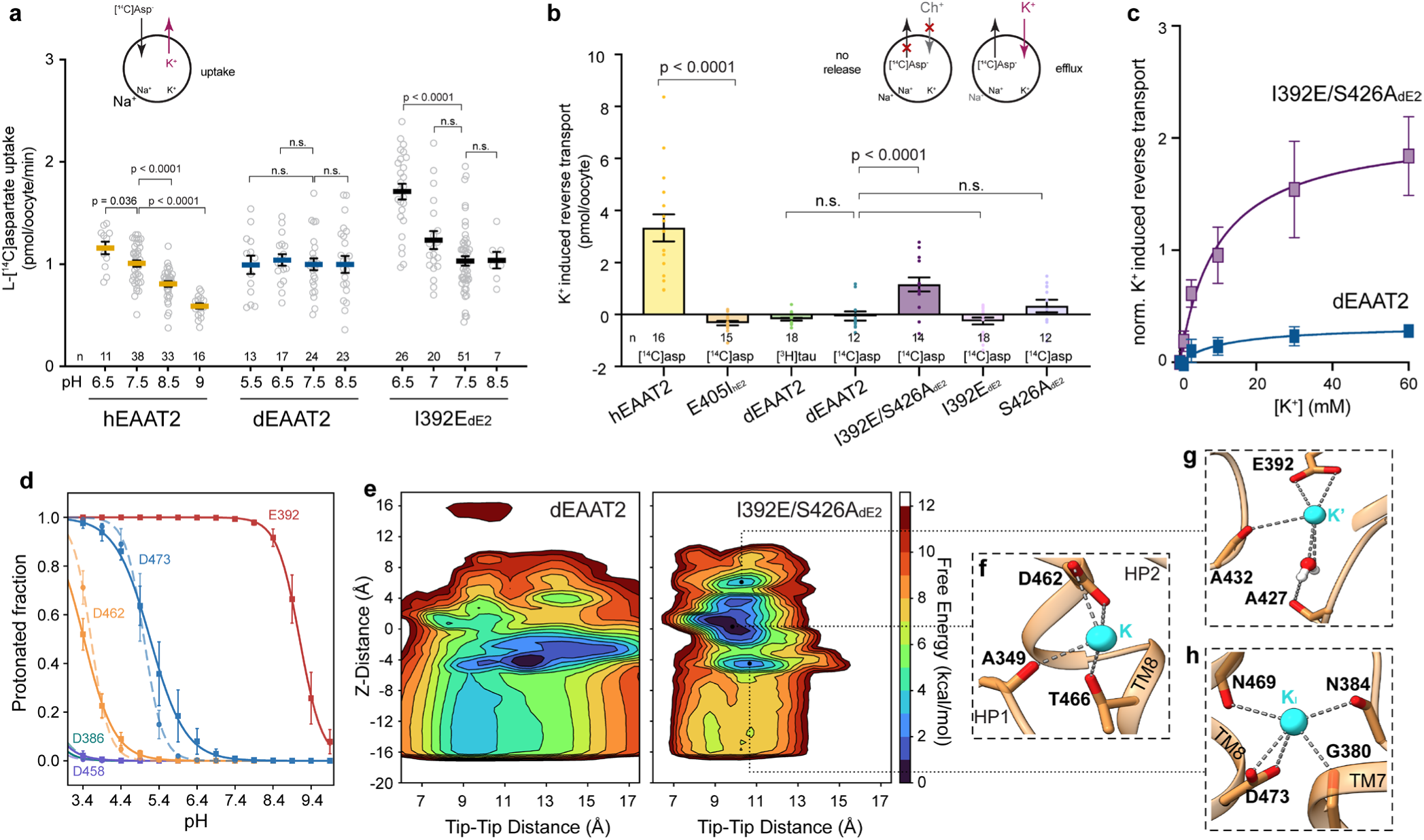
Proton and potassium dependence of hEAAT2 and dEAAT2. (**a**) _L_-[^14^C]aspartate uptake by hEAAT2-, dEAAT2-, and I392E_dE2_-expressing oocytes with various external H^+^ concentrations. Substrate uptake normalised to responses at pH 7.5 from the same batch of oocytes. (**b**) Reverse transport of _L_-[^14^C]aspartate induced by high external K^+^. Background efflux from control oocytes subtracted from each batch of experiments. (**c**) Reverse transport currents in dEAAT2- and I392E/S426A_dE2_-expressing oocytes measured at +60 mV in the presence of 30 μM TBOA with various external K^+^ concentrations (0 – 60 mM). Data was normalised to the currents at [K^+^] = 0 mM, n=5. (**d**) Titration curve from Constant pH Molecular Dynamics (CpHMD) simulations for dEAAT2 and I392E/S426A_dE2_. (**e**) Free energy profiles, projected onto two collective variables (Z-distance and Tip-Tip distance), reveal potential K^+^ binding sites in dEAAT2 (left) and I392E/S426A_dE2_ (right). Tip-Tip distance (TTD) measures the distance (Å) between the C-alpha atoms of residues 348-352 (HP1) and 426-430 (HP2) and the Z-distance measures the Z-coordinate of K^+^ with respect to the mass center of side-chain oxygen atoms of residues D462, T466 (TM8), A349, S350 (HP1), A426, A427, V429 (HP2). Close-up of the three K^+^ binding sites identified in I392E/S426A_dE2_; (**f**) K at a Z-distance of 2-3 Å, (**g**) K’ at a Z-distance of 7 Å, and (**h**) K_I_ at a Z-distance of -4 Å.

Next, we investigated the K^+^ dependence of transport using a radiolabelled substrate efflux assay. Oocytes were preloaded with _L_-[^14^C]aspartate then incubated in a buffer with high (60 mM) K^+^ and low (40 mM) Na^+^ to replicate conditions that would induce reverse transport in the K^+^-dependent transporter hEAAT2^10,42^. Indeed, we observed efflux of preloaded _L_-[^14^C]aspartate in hEAAT2 but not dEAAT2 (Fig. 2b), revealing that the *Drosophila* homologue is K^+^-independent. Several K^+^-binding sites have been proposed in hEAATs^38,40,41,50^, including the double alanine motif in the loop region of HP2 ^43^, which is not conserved in dEAAT2 (Extended Data Fig. 1). To test the role of the double alanine motif in K^+^ coupling, we mutated S426 to alanine in dEAAT2 (S426A_dE2_). K^+^-induced efflux of _L_-[^14^C]aspartate through S426A_dE2_ was not significant, indicating that the double alanine motif alone cannot confer K^+^ dependence (Fig. 3b). Because all human EAATs are coupled to both K^+^ and H^+^, we hypothesised that H^+^ dependence may be required to establish K^+^ dependence. Indeed, the double mutant I392E/S426A_dE2_ demonstrated K^+^-induced _L_-[^14^C]aspartate efflux, whereas the single I392E_dE2_ mutation did not (Fig. 3b). Moreover, abolishing H^+^ dependence in hEAAT2 by mutating E405 (E405I_hE2_) additionally abolished K^+^ dependence (Fig. 3b), rendering the transporter electroneutral (Fig. 1e). To confirm the K^+^ dependence of I392E/S426A_dE2_, we measured reverse transport-activated Cl^-^ currents, which we isolated using the EAAT-specific inhibitor _DL_-TBOA^40^. Unlike WT dEAAT2, currents through I392E/S426A_dE2_ at +60 mV increased as external K^+^ concentration increased (Fig. 3c), indicating successful reconstitution of K^+^ dependence.

To validate these observations, we performed equilibrium and non-equilibrium MD simulations of WT and mutant dEAAT2 in the OF_C_S. In simulations of dEAAT2, I392E_dE2_, and I392E_P_/S426A_dE2_ (I392E_P_ being protonated), the protein remained stable and did not show major structural rearrangements for the length of the simulation (200 ns) (Extended Data Fig. 4a). Substrate also stayed bound during simulations of dEAAT2 and I392E_P_/S426A_dE2_ but the unprotonated I392E_dE2_ variant exhibited an unstable HP2 that allowed substrate to dissociate in all replicas (Extended Data Fig. 4d). This suggests that mutation of I392 destabilises substrate binding, which can be recovered by protonation of E392 (Extended Data Fig. 4b-c). Extensive constant-pH simulations were performed to construct protonation profiles for dEAAT2 and I392E/S426A_dE2_, revealing that the glutamate introduced at position 392 is likely the only permanent protonation site with a pKa value of approximately 9.1 (Fig. 3d), confirming the role of I392E_dE2_ as the H^+^ carrier.

We examined the K^+^-binding ability of I392E/S426A_dE2_ using free-energy calculations on the pre-open Cl^-^-conducting state (_P_ClCS) in the absence of substrate and Na^+^ ions (Fig. 3e; Extended Data Fig. 5). Three potential K^+^-binding sites were identified, with the lowest free-energy minimum site (K site) near the substrate-binding pocket (Fig. 3e; Extended Data Fig. 5c). K^+^ bound at this site was coordinated by residues D462, A349, and T466, resembling the cation/K^+^ binding site in EAAT3^43^ (Fig. 3f). Just above, we identified a secondary, higher free-energy minimum site (K′ site), in which K^+^ interacted with both introduced residues, I392E and S426A (Fig. 3g, Supplementary Video 2). A third minimum was identified towards the intracellular side (K_I_ site), located between Na1 and Na3, in which K^+^ formed interactions with D473, G380, N384, and N469 (Fig. 3h). In WT dEAAT2, K_I_ had the lowest free-energy minimum, consistent with MD simulations on other SLC1A systems^38^. However, the sites resembling K and K′ formed shallow and less stable minima, indicating unfavourable migration of K^+^ to these more extracellular sites in WT transporters (Fig. 3e). The dEAAT2 tip-tip distance profile was also broader, signifying a more flexible and open arrangement of HP1 and HP2, potentially explaining why dEAAT2 acts as an exchanger that only enters the return step of the transport cycle when substrate is bound (Extended Data Fig. 4b). Furthermore, equilibrium MD simulations revealed occasional fluctuations of K^+^ between K and K′ in I392E_P_/S426A_dE2_, in contrast to the stability of K^+^ at K_I_ throughout simulations of K^+^-independent dEAAT2 (Supplementary Video 1, 2). Together, these data reveal that H^+^ dependence is a prerequisite for K^+^ dependence and the I392E/S426A_dE2_ double mutation confers both H^+^ and K^+^ dependence on dEAAT2.

### I392E/S426A_dE2_ also confers Na^+^ dependence

Although all SLC1A family members exhibit a thermodynamically uncoupled Cl^-^ conductance, those dependent on K^+^/H^+^ are electrogenic, whereas K^+^/H^+^-independent members are electroneutral. We therefore wondered whether the I392E/S426A_dE2_ double mutation affects the electroneutrality of dEAAT2. We first confirmed dEAAT2’s electroneutrality by measuring currents in the absence of permeable anions, which abolished currents at all test potentials for WT dEAAT2 as well as the single mutant transporters I392E_dE2_ and S426A_dE2_ (Fig. 4a). In contrast, robust inward currents were observed for hEAAT2 in the same recording conditions due to the transport cycle resulting in net transfer of two positive charges (Fig. 4a). Surprisingly, a similarly inwardly rectifying conductance was observed in the absence of permeable anions for the I392E/S426A_dE2_ double mutant (Fig. 4a), through which transport of _L_-[^14^C]aspartate had become dependent on membrane potential (Fig. 4b). This suggests that the double mutation renders dEAAT2 electrogenic and furthermore that the observed currents are likely due to Na^+^ coupling as K^+^/H^+^ dependence alone would not introduce any net transfer of charge across the membrane.

**Figure 4.**
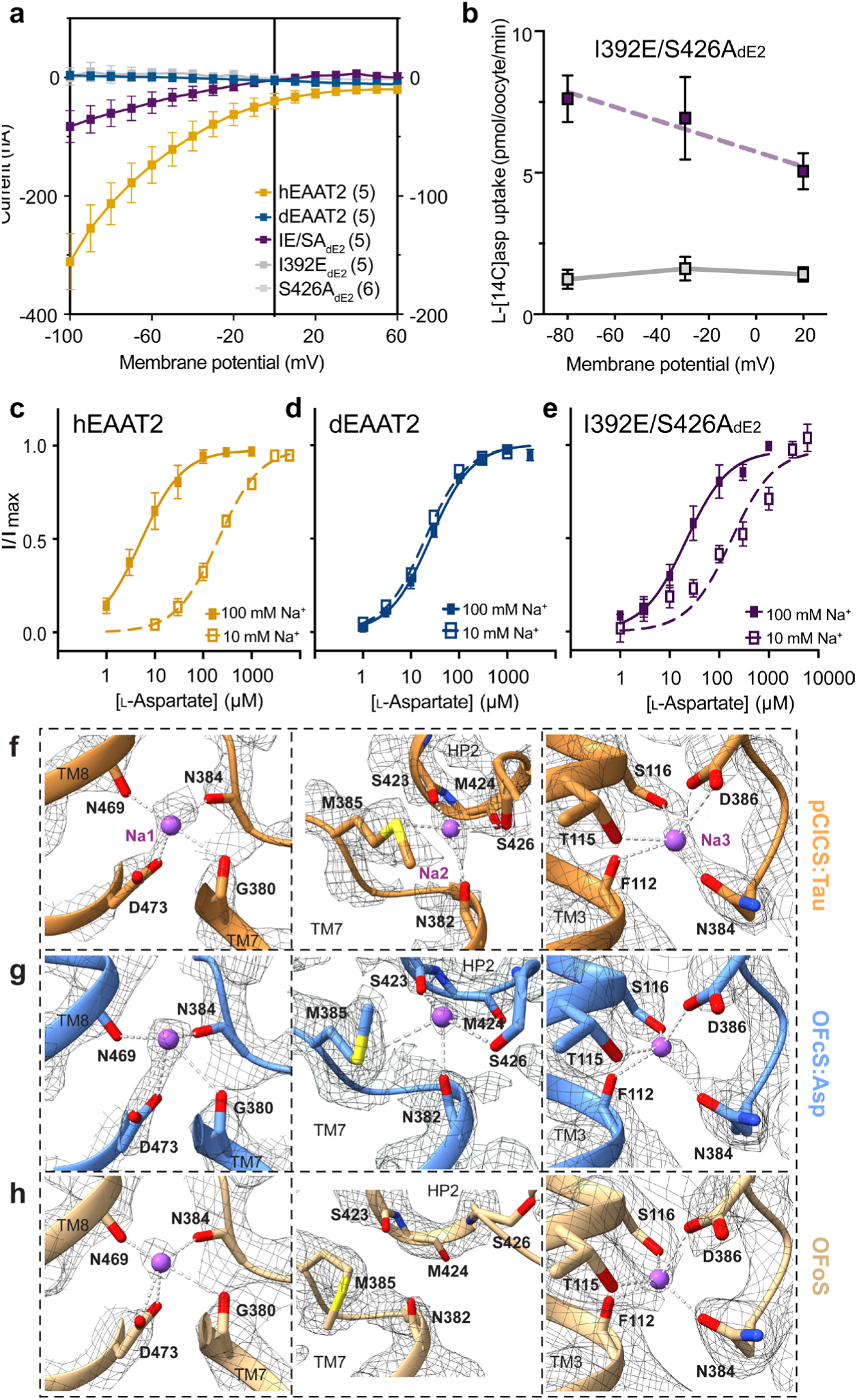
Sodium dependence of dEAAT2. (**a**) IV relationships in response to 100 μM _L_-aspartate in buffers containing non-permeable anions for hEAAT2-expressing oocytes (left y-axis) and dEAAT2 or dEAAT2 mutant-expressing oocytes (right y-axis). (**b**) _L_-[^14^C]aspartate uptake by the double mutant measured at a range of voltages, -80 mV, n=5; -30 mV, n=6; 20 mV, n=7. Background uptake from control oocytes (solid grey line), n=3. Aspartate dose-response curves in the presence of 10 mM Na^+^ (open squares) and 96 mM Na^+^ (filled squares) for (**c**) hEAAT2, (**d**) dEAAT2 and (**e**) I392E/S426A_dE2_ (see Extended Data Table 1 for exact n). Data obtained from at least two different batches of oocytes and presented as mean ± SEM for the indicated number of cells (n) unless otherwise stated. Data analysed using ordinary One-Way ANOVA with Dunnett’s multiple comparison tests by GraphPad Prism 10. The three Na^+^ binding sites; Na1 (left panel), Na2 (middle panel), Na3 (right panel) of (**f**) _P_ClCS:Taurine complex (orange), (**g**) OF_C_S:Aspartate complex (blue), (**h**) OFoS (wheat).

To investigate this Na^+^ dependence, _L_-aspartate dose-response curves were constructed in the presence of either 10 mM or 100 mM external Na^+^. Because WT dEAAT2 is a Na^+^-independent exchanger, the aspartate and taurine dose-response curves in 10 mM and 100 mM Na^+^ were indistinguishable (Fig. 4d; Extended Data Fig. 6i). In contrast, the dose-response curve for the Na^+^-coupled transporter hEAAT2 displayed a rightward shift following the ∼10-fold increase in Na^+^ concentration (Fig. 4c). Moreover, although neither of the single mutants demonstrated any change when Na^+^ was increased (Extended Data Table 1), the I392E/S426A_dE2_ double mutant showed a clear dependence on external Na^+^ concentration (Fig. 4e) such that the apparent affinity for aspartate decreased by ∼8-fold following a ∼10-fold reduction in Na^+^ concentration (Extended Data Table 1). Together, these findings demonstrate that the I392E/S426A_dE2_ double mutation converted dEAAT2 from an electroneutral exchanger to a Na^+^/H^+^/K^+^-dependent electrogenic transporter, resembling its human counterpart.

The ability of the I392E/S426A_dE2_ double mutation to confer Na^+^ dependence on dEAAT2 was intriguing. Because the amino acid residues that form the Na1-3 sites are conserved in dEAAT2 (Extended Data Fig.1), we determined high-resolution cryo-EM structures of dEAAT2 with and without Na^+^ to investigate the involvement of these sites in substrate transport. In the presence of saturating concentrations of Na^+^ and taurine, we observed clear densities for Na^+^ at all three Na^+^-binding sites in _P_ClCS (Fig. 4f). In the presence of saturating concentrations of Na^+^ and aspartate, we also observed non-protein densities at Na1-3 in the OF_C_S (Fig. 4g; Extended Data Fig. 3c,d; Supplementary Table 1).

We sought to further investigate the role of Na^+^ by determining the structure of dEAAT2 in the absence of Na^+^ and substrate, but the protein was too unstable to resolve any structures. However, we were able to determine open state structures of dEAAT2 in an OFS (OF_O_S) and IFS (IF_O_S) by reducing the concentration of Na^+^ and aspartate (Extended Data Fig. 3c,d; Supplementary Data Fig. 7-8; Supplementary Table 1). These structures revealed a transition of residues 93–103 from the start of TM3 in the OF_O_S to the end of TM2 in the IF_O_S (Supplementary Data Fig. 9), relaxing the intracellular hinge between the scaffold and transport domains, and allowing the transport domain to lower as in other elevator transporters^31,34,43^. We did not observe density for Na^+^ at the Na2 site in either of the open structures (Fig. 4h and Extended Data Fig. 6d-f), consistent with Na2 being disrupted when HP2 is in an open conformation in other SLC1A family members^34,43,50^. Indeed, MD simulations of dEAAT2 showed that the Na^+^ ion at Na2 frequently unbound, increasing the distance between the tips of hairpin 1 (HP1) and HP2 to 14 Å, thus allowing the substrate to unbind (Extended Data Fig. 4).

Because HP2 is closed in the _P_ClCS:taurine complex (Fig. 4f), Na2 can be coordinated by the main chain oxygen atoms from S426 (HP2), S423 (HP2), N382 (TM7), and the side chain of M385 (TM7). However, in the OF_C_S:aspartate complex, where HP2 is also closed, M385 adopts a different rotamer, disengaging from the Na2 network (Fig. 4g) and likely becoming more flexible (Extended Data Fig. 6g,h). M385 is part of the TM7 “NMD” motif, thought to be essential for Na^+^ coupling in SLC1A members^10,34,46^. The ability of HP2 to close and initiate substrate transport via dEAAT2 while having lost one of the two Na^+^-TM7 interactions may explain why dEAAT2 function is less dependent on Na^+^. Furthermore, previous studies have shown that the side chains of residues equivalent to M385 in other SLC1A Na^+^-dependent transporters swing away from the substrate-binding site in their open state^33,35,51^, disengaging HP2 and TM7 to enable substrate release. However, this reorientation is less significant in the open conformations observed in this study, where M385 is in a similar position in the presence or absence of Na^+^ and substrate (Extended Data Fig. 6g,h). These data demonstrate that dEAAT2 is a Na^+^ independent exchanger and the I392E/S426A_dE2_ double mutation confers Na^+^ dependence on substrate transport by functionally enabling the three canonical Na^+^-binding sites.

Given the importance of E392_dE2_ and A426_dE2_ for ion-coupled transport, we performed ancestral protein reconstruction to understand the evolutionary origin of these changes, and whether the ion-independent exchange function of dEAAT2 emerged from ion coupling function, or vice versa^52^. Alignment of SLC1A sequences from Bilateria were used to generate a maximum-likelihood phylogeny and reconstruct ancestral protein sequences, showing stepwise sequence changes in SLC1A homologs from ancient animals to modern arthropods and mammals (Extended Data Fig. 7a). The most plausible phylogeny suggests that an ancestral bilaterian SLC1A transporter, Bilat_ANC_, possessed the E392/A426 residues required for ion-coupled transport. This is consistent with ion-coupled SLC1A function prevailing throughout all kingdoms of life^51,53,54^. An apparent gene duplication event resulted in ancestral precursors of dEAAT1/hEAAT1,3,4,5 (Ion1_ANC_) and dEAAT2/hEAAT2 (Ion2_ANC_), both with E392/A426. These residues are unaltered in Ion1_ANC_ descendants; however, signatures of ion-independent electroneutral exchange emerged sequentially in an Ion2_ANC_ lineage unique to protostomes (Extended Data Fig. 7b). E392_dE2_ was first substituted with a hydrophobic residue (Exc1_ANC_), which our data in hEAAT2 suggests would result in loss of ion coupling, regardless of A426_dE2_ status (Figure 3b). The subsequent replacement of A426_dE2_ with serine may have occurred as the double alanine motif in HP2 was no longer required to facilitate ion coupling.

## Discussion

We have shown that the *Drosophila* amino acid transporter, dEAAT2, utilises a distinct substrate transport mechanism from hEAATs, despite having high amino acid sequence identity. dEAAT2 is a Na^+^/H^+^/K^+^-independent electroneutral exchanger that catalyses aspartate and taurine exchange across the membrane while activating an uncoupled Cl^-^ conductance. Taurine transport via dEAAT2 requires the presence of Cl^-^, whereas aspartate transport does not, which we propose is due to taurine lacking a second carboxylate and therefore requiring a Cl^-^ ion to occupy the same substrate-binding pocket as aspartate. Intriguingly, unlike in other eukaryotic SLC1A members, where the IFS is the most favourable state^43,50,55^, dEAAT2 prefers a previously unreported intermediate state that is close to the open Cl^-^ conducting state (ClCS) observed in the prokaryotic homologue Glt_Ph_^15^. The combination of the unique lack of ion dependence of dEAAT2 and its propensity to exist in the _P_ClCS allowed us to interrogate SLC1A ion coupling mechanisms and probe K^+^ interactions on the return step of the transport cycle due to the likelihood that the K^+^-bound IFS is a high-energy state^43,50^.

The surprising lack of dependence on external Na^+^ for substrate transport in dEAAT2 is in sharp contrast to all other eukaryotic SLC1A family members characterised thus far. The Na^+^-binding sites are mostly conserved in dEAAT2^31,34,56,57^ and our results suggest that Na^+^ can still bind to dEAAT2 even though it is not essential for transporter function. This is in contrast to an earlier study suggesting that taurine transport via dEAAT2 is Na^+^-coupled^22^. However, these earlier results were likely confounded by the effect of Cl^-^ removal on taurine transport, as mannitol was used as an osmotic replacement for both Na^+^ and Cl^-^. The lack of Na^+^ dependence in dEAAT2 could be due to the loss of M385_dE2_ coordination, which is critical for Na2 binding, limiting flexibility of the hinge between TM7a and TM7b in the vicinity of the Na^+^-binding sites compared to other SLC1A members. A similar phenomenon was observed in the electroneutral exchanger hASCT2^58^. We speculate that exchangers in the SLC1A family may have a more rigid TM7 that maintains M385_dE2_ in a similar orientation in both open and closed states of HP2, enabling Na2 to be more readily available for Na^+^ rebinding when HP2 closes, thus reducing dependence on external Na^+^.

E405_hE2_ has been proposed to be the H^+^ carrier in the SLC1A family^33^ and to play a role in K^+^ coupling^41,48^. dEAAT2, which possesses an isoleucine at the equivalent position, displays no dependence on Na^+^, H^+^, or K^+^; however, the introduction of a glutamate residue in the *Drosophila* exchanger (I392E_dE2_) introduces H^+^ dependence, but not Na^+^ or K^+^ dependence, even though the Na^+^-binding sites are conserved in dEAAT2. Only when I392E_dE2_ is combined with a second mutation in HP2, S426A_dE2_, is K^+^ and Na^+^ dependence introduced into the *Drosophila* homologue. We suspect the introduction of Na^+^ dependence to I392E/S426A_dE2_ may be linked to a requirement for Na^+^ binding to destabilise a salt bridge between the H^+^ carrier E405_hE2_ and a highly conserved substrate-coordinating residue R478_hE2_ so the substrate-binding site can be established^33,35,59^. K^+^ dependence is likely due to deprotonation of E405_hE2_ in the inward-facing state being necessary to allow K^+^ access to its binding site^43^. Thus, K^+^ dependence was introduced only when dEAAT2 became H^+^-dependent, and Na^+^ became important for dEAAT2 function only when H^+^/K^+^ dependence was present. Our phylogenetic analyses demonstrate that ion-independent electroneutral exchange emerged from ion-coupled transport, exclusively in protostomes. The initial I392E_dE2_ substitution suggests that both K^+^/H^+^ dependence were simultaneously eliminated, while some transporters, including dEAAT2, subsequently gained the A426S_dE2_ variant. As the residues that confer taurine specificity are unknown, it is unclear in which order ion-independent electroneutral exchange and taurine specificity emerged. Interestingly, following the I392E_dE2_ substitution, molluscs diverged to contain a bulkier hydrophobic residue at the 426_dE2_ position, potentially altering substrate specificity.

Together, these findings have enabled the dissection of the molecular mechanisms underlying ion coupling in SLC1A transporters, revealing that dEAAT2 is a powerful tool to investigate transport mechanisms in vertebrate members of the family. Specifically, we identified the molecular basis of K^+^ dependence and how this is linked to both H^+^ and Na^+^ dependence. Further, our finding that two mutations are sufficient to convert dEAAT2 from an ion-independent electroneutral exchanger into a Na^+^/H^+^/K^+^-dependent electrogenic transporter reveals how high-order ion-coupling mechanisms are achieved in SLC1A transporters (Extended Data Fig. 8). Our data also explain previous challenges in functionally verifying these ion-binding sites, which were likely due to the knockout of one binding site impacting the function of the others^38–41^. These findings thus increase our understanding of the intricacies of ion coupling and provide insight into the molecular differences between transport and exchange of substrates in the SLC1A family. Crucially, our insight could inform the design of novel therapeutics to target the K^+^-binding step of the transport cycle, which would prevent excitotoxicity caused by reverse transport in disease states such as ischaemic stroke^60,61^.

## Methods

### Molecular biology, oocyte harvesting, and preparation

Complementary DNA (cDNA) encoding EAAT2 from *Drosophila melanogaster* (dEAAT2) was obtained from Professor Donald van Meyel, McGill University and subcloned into the plasmid oocyte transcription vector (pOTV) for expression in *Xenopus laevis* oocytes. Site-directed mutagenesis was performed with the Q5 Site-directed Mutagenesis Kit (New England Biolabs, Inc). Primers encoding the desired mutations were designed via the online NEBaseChanger tool (New England Biolabs, Inc) and synthesised by Sigma-Aldrich (Sydney, Australia). DNA products were transformed into NEB® 5-alpha competent *E. coli* cells and plasmid DNA was purified with PureLink® Quick Plasmid Miniprep Kit (Invitrogen). Sequence-confirmed WT and mutant cDNAs were linearized with *Spe1* restriction enzyme (New England BioLabs), prior to transcription into cRNA by T7 RNA polymerase (mMESSAGE mMACHINE® Kit, Ambion, USA). Constructs were sequence-confirmed using Big Dye Terminator (BDT) labelling reaction on both strands by the Australian Genome Research Facility (Sydney, Australia).

*Xenopus laevis* frogs were purchased and imported from NASCO International (Wisconsin, USA). Oocytes were surgically removed from anaesthetised *Xenopus laevis* frogs after the administration of tricaine for 12 minutes. Surgical procedures were approved by the University of Sydney Animal Ethics Committee. Oocytes were defolliculated with 2 mg/mL collagenase for 1 hour and 23 nL of cRNAs encoding desired transporters were injected into stage V defolliculated oocytes, which were stored at 18 °C in Storage Solution (96 mM NaCl, 2 mM KCl, 1 mM MgCl_2_, 1.8 mM CaCl_2_, 2.5 mM sodium pyruvate, 0.5 mM theophylline, 50 µg/mL gentamycin and 50 µg/mL tetracycline, 5 mM hemi-Na^+^ HEPES, pH 7.5) for at least 48 hours to allow sufficient protein expression on cell membrane.

### Electrophysiology

Between 2 and 4 days after injections, currents were recorded using the two-electrode voltage clamp technique with a Geneclamp 500 amplifier (Axon Instruments) interfaced with a PowerLab 2/20 chart recorder (ADInstruments) and a Digidata 1322A (Axon Instruments), which was used in conjunction with Chart software (ADInstuments; Axon instruments). All recordings were made with a bath grounded via a 3 M KCl/1% agar bridge linking to a 3 M KCl reservoir containing a Ag/AgCl_2_ ground electrode to minimize offset potentials. Current–voltage (IV) relationships for substrate-elicited conductance were determined by measuring substrate-elicited currents during 245-ms voltage pulses between −100 mV and +60 mV at 10 mV steps. Background currents were eliminated by subtracting currents in the absence of substrate from substrate-elicited currents at corresponding membrane potentials. Experiments determining half saturations (*Km*) of various substrates were conducted in standard frog Ringer’s solution (ND96: 96 mM NaCl, 2 mM KCl, 1 mM MgCl_2_, 1.8 mM CaCl_2_, 5 mM hemi-Na^+^ HEPES, pH 7.5) with an increasing dose of taurine, _L_-aspartate, and _L_-glutamate, unless otherwise stated. Substrate-elicited currents (I) at -60mV were fitted to the Michaelis-Menten equation by least squares:

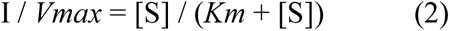

where *Km* is the substrate concentration required to reach half-maximum response, *Vmax* is the maximum response, and [S] is the substrate concentration. Reverse transport currents were isolated by application of 100 μM of the EAAT selective inhibitor D,L-threo-betahydroxyaspartate (TBOA) in KNO_3_ buffers ([K^+^] from 0 – 60 mM). The current recorded at +60 mV at each K^+^ concentration in the presence of TBOA was normalised to the current recorded at [K^+^] = 0 mM and plotted on a logarithmic scale.

### Radiolabelled uptake and efflux in oocytes

Forward transport (uptake) was assayed by incubating oocytes expressing the various transporters and control (uninjected) oocytes with 10 µM _L_-[^14^C]aspartate, [^3^H]taurine or _L_-[^3^H]glutamate for 10 minutes; uptake was terminated by washing oocytes three times with ice-cold ND96. Uptake under voltage clamp was measured by clamping oocytes at a range of membrane potentials from -80 mV to 0 mV and incubating them with 30 μM _L_-[^14^C]aspartate or [^3^H]taurine under indicated membrane potential for 2 minutes, followed by a 5-minute washout period. For the efflux assay, oocytes were preloaded with radiolabelled substrate by direct injection or by incubating oocytes with 10 μM _L_-[^14^C]aspartate or [^3^H]taurine for 30 minutes. Preloading was terminated by three washes with ice-cold ND96, followed by three washes with choline buffer at room temperature. Efflux was initiated by placing the preloaded oocytes into a reverse transport buffer (40 mM NaCl, 60 mM KCl, 1 mM MgCl_2_, 1.8 mM CaCl_2_, 5 mM HEPES, pH7.5, pH with TrisBase), an exchange buffer (ND96 containing 300 μM unlabelled substrates), or a control buffer (96 mM ChCl, 2 mM KCl, 1 mM MgCl_2_, 1.8 mM CaCl_2_, 5 mM HEPES, pH7.5, pH with TrisBase choline buffer), and incubated at room temperature for 20 minutes with gentle agitation.

Following both uptake and efflux assays, oocytes were lysed with 1M NaOH and 1% SDS before the addition of scintillant (Optiphase HisSafe, PerkinElmer). Both the lysed oocytes and the efflux solution were sampled for counting using a MicroBeta TriLux scintillation counter (PerkinElmer). Uptake was normalised to uptake under a standard condition (ND96) of each individual experiment unless otherwise stated. For anion experiments, 96 mM NaCl was substituted with 96 mM anion sodium salts (NaMES, NaF, NaBr, NaI and NaNO_3_). For Cl^-^ titration experiments, NaMES was used as a substituent to maintain equivalent osmolarity across buffers with different Cl^-^ concentrations, from 0 mM (150 mM NaMES buffer) to 150 mM; at Na^+^-free conditions, choline chloride (ChCl) was used to provide Cl^-^ with NMDG^+^ and MES^-^ used as Na^+^ and Cl^-^ substituents, respectively. Reverse transport or efflux is represented as the total amount radioabelled substrate (in pmols) effluxed from the oocyte into the external solution over 30 minutes.

### Aspartate competition assay

Increasing concentrations of aspartate- and taurine-related compounds were applied up to 1 mM (3 mM for β-alanine and glutamate) to dEAAT2 in the presence and absence of Cl^-^. Dose-dependent inhibition curves were obtained if compounds were able to bind and inhibit aspartate uptake, analysed using the following equation:

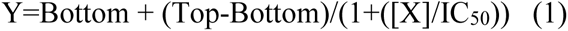

where Y is the normalised _L_-[^14^C]aspartate uptake under each condition, top and bottom are the maximum or minimum responses, [X] is the concentration of compounds applied, and IC_50_ is the concentration of the compound that gives a response halfway between Bottom and Top.

### Protein expression and purification

Wild-type *Drosophila melanogaster* EAAT2 (dEAAT2) was cloned into a pFastBac vector containing a TEV protease cleavage site at the C-terminus, followed by a green fluorescent protein (GFP) and an 8×His tag. Recombinant proteins were expressed in *Spodoptera frugiperda* Sf9 insect cells (Invitrogen) via baculovirus-mediated infection. Sf9 cells were cultured in Insect-XPRESS^TM^ protein-free medium (Lonza) to a density of 4 × 10^6^ cells/mL and infected with baculovirus encoding either wild-type or mutant dEAAT2. Infected cultures were incubated at 27 °C and harvested by centrifugation 48 h post-infection. Cell pellets were resuspended and homogenized in buffer containing 50 mM Tris-HCl (pH 8.0), 500 mM NaCl, 1 mM TCEP-HCl, 2% (w/v) dodecyl-β-D-maltopyranoside (DDM; Anatrace), 0.4% (w/v) cholesteryl hemisuccinate (CHS; Anatrace), 10% (v/v) glycerol, 1 mM MgCl₂, and 10 mM _L_-aspartic acid sodium salt. DNase I, Benzonase® Nuclease, and cOmplete™ Protease Inhibitor Cocktail (all from Merck) were also included.

The homogenate was stirred at 4 °C for 1 h, followed by centrifugation at 40,000 × *g* for 30 min. The supernatant was filtered using a glass fibre prefilter and incubated with Ni-NTA resin (Qiagen) for 1 h at 4 °C. Resin was then packed into a glass column and washed with 20 column volumes of buffer containing 50 mM Tris-HCl (pH 8.0), 500 mM NaCl, 1 mM TCEP-HCl, 50 mM imidazole, 0.06% glyco-diosgenin (GDN), 5 mM _L_-aspartic acid sodium salt, and 5% glycerol. Protein was eluted using a buffer of identical composition, except with 200 mM NaCl and 300 mM imidazole. The GFP and His tag were removed by overnight incubation with TEV protease (Merck; 10 U/mg protein) at 4 °C. The protein was further purified by size-exclusion chromatography (SEC) using a Superose 6 Increase 10/300 GL column (Cytiva). For _L_-aspartate-bound samples, the SEC buffer comprised 20 mM Tris-HCl (pH 8.0), 200 mM NaCl, 1 mM TCEP-HCl, 0.01% GDN, and either 1 mM or 5 mM L-aspartic acid sodium salt. For taurine-bound samples, 10 mM taurine replaced _L_-aspartate in the membrane resuspension, wash, and elution buffers. The corresponding SEC buffer contained 20 mM Tris-HCl (pH 8.0), 200 mM NaCl, 1 mM TCEP-HCl, 0.01% GDN, and 5 mM taurine.

### Cryo-EM data acquisition and processing

3 μL of freshly purified dEAAT2 protein at ∼6 mg/mL was placed on the glow-discharged C-flat™ holey carbon grids (MiTeGen; Cu R1.2/1.3) and blotted for 5 s and flash-frozen in liquid ethane cooled by liquid nitrogen using a Vitrobot Mark IV (Thermo Fisher Scientific Inc.). The grids were clipped and subsequently transferred to a FEI Titan Krios (300 kV, Gatan K3 equipped post a BioQuantum energy filter) cryo-electron microscope. Automated data collection was performed using aberration-free image shift (AFIS) as implemented in EPU 2 software (Thermo Fisher Scientific). Images were collected with a defocus range of ∼ –0.4 to –1.4 μm. The microscope was operated at a nominal magnification of 105,000 x in EFTEM nanoprobe mode with a physical pixel size of 0.82 Å. The total dose applied was 60 e^−^/Å^2^ at a dose rate of 9.515 e^−^/pixel/second. The energy filter was operated with a slit width of 10 eV, the Gatan K3 camera was operated in correlated double sampling (CDS) mode, and movies were saved as TIFF files with 60 subframes. The beam-induced motion within each dose-fractionated cryoEM micrograph was corrected using MotionCor2^62^. The defocus and astigmatism values of the micrographs were determined using CTFFIND4^63^. Particles were picked using Topaz^64^. The single particle data analysis was performed using RELION4^65^ and cryoSPARC4^66^ following customized workflows (Supplementary Fig. 3, 6, and 8). It is worth noting that different data processing strategies were initially tested, which helped to understand the nature of the data sets and design the final data processing strategies.

For the dEAAT2 sample prepared in the presence of 5 mM taurine, a total of 6,692 cryo-EM micrographs were collected. Of these, 5,812 were selected for further processing based on contrast transfer function (CTF) estimation using CTFFIND4. Particle picking using Topaz yielded 2,811,414 particles (binned 1.875×; 128 × 128 pixels; 1.5375 Å/pixel), which were subjected to two rounds of 2D classification. A subset of 492,879 particles from the best 2D classes was selected for 3D classification in RELION 4.0 with C3 symmetry. An initial model was generated de novo with C3 symmetry. From three 3D classes, the best class containing 378,973 particles was selected, re-extracted at full pixel size (240 × 240 pixels; unbinned, 0.82 Å/pixel), and subjected to 3D auto-refinement, particle polishing, 2D classification, and CTF refinement, resulting in a 2.94 Å reconstruction with C3 symmetry (Supplementary Fig. 3). To further improve the resolution that was limited by the imperfect symmetry of the dEAAT2 trimer, symmetry expansion and focused refinement were performed. The particles from the 3D auto-refinement were symmetry-expanded with the C3 symmetry using *relion_particle_symmetry_expand*. The expanded particles were subjected to focused 3D classification, and a subset of 314,902 asymmetric units was selected for subsequent 3D auto-refinement, particle polishing, and CTF refinement, yielding a reconstruction at 2.46 Å resolution. Following deduplication, 211,799 particles were transferred to cryoSPARC and refined using non-uniform refinement with C3 symmetry, resulting in a final global reconstruction at 2.38 Å resolution. Both the trimeric and protomeric reconstructions revealed an identical chloride-conducting pre-open state (_P_ClCS). To avoid the mismatch of size and shape between the reference of the small protomer and the particle images of the large dimer during 3D classification and 3D auto-refinement, a special masking method with composite masks was used in a modified version of RELION4 as follows^67,68^. In addition to a regular reference mask covering a single protomer (yellow mask in Supplementary Fig. 3), a secondary mask (grey mask in Supplementary Fig. 3) encompassing the adjacent protomer and surrounding detergent micelle was applied. During 3D classification and 3D auto-refinement, this secondary mask was low-pass filtered to 20 Å. This composite masking approach preserved low-resolution information from the full particle to guide global alignment, while allowing high-resolution features within the masked protomer to dominate classification and refinement, thus improving local structural detail and alignment accuracy.

For the dEAAT2 sample prepared with 5 mM _L_-aspartate, a total of 6,644 cryo-EM micrographs were collected. Of these, 5,776 were selected for further processing based on CTF estimation using CTFFIND4. Particle picking with Topaz yielded 2,347,765 particles (binned 1.875×; 128 × 128 pixels; 1.5375 Å/pixel), which were subjected to 2D classification. A subset of 723,542 particles from the best 2D classes was selected for 3D classification in RELION 4.0 using C3 symmetry. An initial model was generated de novo with C3 symmetry, and the best class, comprising 467,981 particles, was selected from three 3D classes. These particles were re-extracted at full pixel size (240 × 240 pixels; unbinned, 0.82 Å/pixel) and subjected to 3D auto-refinement with C3 symmetry, yielding a reconstruction at 3.51 Å resolution. The refined particles were then symmetry-expanded and subjected to focused 3D classification using a composite masking strategy, as described above. From ten resulting classes, three distinct conformational states were identified: outward-facing closed, Cl⁻-conducting pre-open state (_P_ClCS), and inward-facing open. For the _P_ClCS bound with aspartate, 121,695 particles were selected and subjected to 3D auto-refinement, CTF refinement, and particle polishing, yielding a final reconstruction at 2.94 Å. For the outward-facing closed state, 159,155 particles underwent the same refinement workflow, resulting in a 2.77 Å reconstruction. For the inward-facing open state, 324,707 particles were further subjected to focused 3D classification, and a subset of 102,137 particles was selected. These were refined through 3D auto-refinement, particle polishing, 2D classification, and CTF refinement, ultimately achieving a reconstruction at 3.22 Å resolution (Supplementary Fig. 6).

For the dEAAT2 sample prepared with 1 mM L-aspartate, a total of 7,296 cryo-EM micrographs were collected. Of these, 6,554 were selected for further processing based on CTF estimation using CTFFIND4. Particle picking was performed using Topaz, yielding 1,971,087 particles (binned 1.875×; 128 × 128 pixels; 1.5375 Å/pixel), which were subjected to 2D classification. From the best 2D classes, 342,854 particles were selected for 3D classification in RELION 4.0, using C3 symmetry. An initial model was generated de novo with C3 symmetry, and the best 3D class (201,160 particles) out of three was selected for further processing. These particles were re-extracted at full pixel size (240 × 240 pixels; unbinned, 0.82 Å/pixel) and refined using 3D auto-refinement with C3 symmetry, yielding a 3.5 Å reconstruction. Subsequent 3D classification with relaxed C3 symmetry revealed a hybrid trimer, with one monomer in the outward-facing open state and two subunits in the inward-facing open state. This class, consisting of 75,300 particles, was refined further using 3D auto-refinement and CTF refinement with relaxed C3 symmetry, resulting in a reconstruction at 3.2 Å resolution. Particles were then subjected to Bayesian polishing and transferred to cryoSPARC for further refinement. A 2D classification step retained 74,968 high-quality particles, which were refined using non-uniform refinement to achieve a final global resolution of 2.64 Å. Local refinement with a soft mask encompassing a single protomer yielded reconstructions at 2.66 Å for the outward-facing open monomer and 2.67 Å for the inward-facing monomer (Supplementary Fig. 8).

### Model building and refinement

The initial models for protomers were built de novo from the cryo-EM density and used as initial templates for trimeric models. The models were adjusted into the density using Coot^69^ and ISOLDE^70^, and the final models were iteratively real-space refined and validated in PHENIX^71,72^ using default parameters. All structure figures were made with ChimeraX^73^.

### Membrane embedding and equilibrium MD simulations

The simulation system was designed based on the experimental model of the dEAAT2 OF_C_S and _P_ClCS. Using the Membrane Builder^74–76^ module of CHARMM-GUI^77^, the OFS structure was embedded in a pre-equilibrated patch of glia-like membrane containing 1-palmitoyl-2-oleoyl-sn-glycero-3-phosphocholine (POPC), 1-palmitoyl-2-oleoyl-sn-glycero-3-phospho-ethanolamine (POPE), 1-palmitoyl-2-oleoyl-sn-glycero-3-phosphoserine (POPS), cholesterol (Chol), and a small quantity of 1-palmitoyl-2-oleoyl-sn-glycero-3-phosphoinositol (POPI), POPI24, and POPI33 lipids^4,78^. An asymmetric bilayer was constructed with Chol:POPC:POPE:POPS:POPI:POPI24:POPI33 at 46:30:4:17:1:1:1 in the cytoplasmic leaflet and Chol:POPC:POPE at 45:48:7 in the extracellular leaflet. The system was solvated with TIP3P water^79^ and neutralized with Na^+^ and Cl^−^ ions to 150 mM salt concentration. All simulations were performed in NAMD 3.0^80^ with the CHARMM36m^81,82^ force fields. Temperature was maintained at 310 K using a Langevin thermostat, and pressure was held at 1 bar using the Nose-Hoover Langevin piston method^83,84^. Periodic boundary conditions were applied, long-range electrostatics interaction were computed using the Particle Mesh Ewald method^85^, and a 2 fs integration time step was used. All titratable residues were kept in their standard protonation states. The system was energy-minimized and equilibrated prior to production. During equilibration, protein, membrane, and ligands (substrate and three sodium ions) harmonically restrained to their initial positions with a spring constant of 5 kcal mol^-1^ Å^-2^. The restraints were gradually released during 3 ns and 5 ns equilibrium simulations for the membrane and for the protein and ligands, respectively, followed by a 200 ns unconstrained equilibrium simulation. Configurations from the trajectories of the unconstrained equilibrium simulation were saved at 100 ps intervals. All visualization and analysis were performed using VMD^86^ together with in-house scripts.

### pKa calculations with constant-pH MD simulations

While equilibrium MD suggested that E392 could serve as a proton-binding site, we further validated this through extensive constant-pH (cpH) MD simulations. In this approach, selected protonatable sites can dynamically change their protonation states during the simulation to explore lower-energy configurations. We examined putative protonation sites in WT and in the I392E-S426A variant of the OF conformation in the presence of Na^+^ and substrate, to determine the protonation profiles of luminal residues near the binding site. Four sites were considered in WT (D386, D458, D462, D473) and five in I392E/S426A_dE2_ (the same four plus E392), chosen based on pKa predictions by PROPKA^87^ and prior reports in similar glutamate transporters^33,43,48,88^. Constant-pH simulations were carried out using the hybrid nonequilibrium MD/Monte Carlo (neMD/MC) algorithm implemented in NAMD^89^. Assuming no long-range allosteric coupling between protomers, equivalent sites in each protomer were treated as independent replicas, while all other residues were kept at their default ionization states. The results from these simulations were combined using the binless WHAM procedure^89,90^. Each constant-pH run comprised 3000-3500 neMD/MC cycles with 50 ps nonequilibrium switches followed by 10 ps equilibrium MD (an upper limit of 180-200 ns). For WT, 13 pH values were simulated (3.49.4 in 0.5-unit increments), and for I392E/S426A_dE2_, 14 pH values were simulated (3.4-9.9 in 0.5-unit increments). To maintain charge neutrality, a Galvani offset potential^91^ was applied and calibrated by measuring the pKa of an additional aspartic dipeptide embedded and restrained in bulk solvent. Substrate and Na^+^ ions were also slightly restrained with a spring constant 5 kcal mol^-1^ Å^-2^ to prevent unbinding during cpH simulations. Convergence of the constant-pH calculations was assessed by monitoring the estimated pKa values of the investigated residues as a function of cumulative simulation time. Equivalent sites in the three protomers were treated as independent replicas, as described above, and their combined estimates were monitored throughout the simulations (Supplementary Fig. 10a,b).

### Free energy calculations to investigate K^+^ binding sites

To investigate the most favourable K^+^ binding site and calculate the free energy associated with different sites, we used the on-the-fly probability enhanced sampling (OPES) method^92–94^. OPES is a nonequilibrium MD approach, developed as an advancement of metadynamics^95,96^ and has been shown to offer significant improvements in efficiency and accuracy^92–94,97,98^. Like metadynamics, OPES estimates free-energy landscapes (or, more precisely, the potential of mean force) as a function of carefully chosen collective variables that capture the underlying physics of the process. OPES is implemented in the Collective Variables (Colvars) module^99^ with a multi-walker option available in both metadynamics and explore modes. We chose the multi-walker OPES scheme for its efficiency and its ability to assess convergence of sampling. OPES simulations were performed for dEAAT2 and I392E/S426A_dE2_ in the _P_ClCS, both in the absence of ligands (substrate and Na^+^ ions). The collective variables were: (i) the Tip-Tip distance, defined as the distance between the Cα atoms of residues 348-352 (HP1) and 426-430 (HP2), and (ii) the Z-coordinate of K^+^ with respect to the centre of mass of side-chain oxygen atoms of residues interacting with the substrate, namely D462 and T466 (TM8), A349 and S350 (HP1), and A426, A427, and V429 (HP2). Additionally, K^+^ was restrained within a cylinder of radius 12 Å and height 36 Å, aligned with the membrane normal and centred on the substrate-binding site. Two-dimensional free-energy profiles were reconstructed by reweighting the trajectories using the applied biasing energy. Convergence of the OPES calculations was assessed by monitoring the self-convergence of the reconstructed two-dimensional free-energy landscapes. At cumulative simulation time t, the free-energy landscape was compared with the final free-energy estimate by calculating the root-mean-square deviation (RMSD) over the sampled collective-variable space. The simulations were considered self-converged when the RMSD reached and remained below *k_B_T* (Supplementary Fig. 10c).

For the OPES simulations, the multiple-walker^99^ option was used with 16 walkers in metadynamics mode^92–94^. To obtain initial conformations, Replica Exchange with Solute Tempering (REST2)^100^ simulations were first performed. In these runs, all atoms in the binding-site region and the K^+^ ion in each protomer were defined as solute atoms for scaling. Twenty-four replicas were used, spanning effective temperatures of 310-620 K with an exchange attempt frequency of 0.2 ps^-1^. Each replica was simulated for 40 ns. Through these simulations, we noticed that two important measures for K^+^ binding are the Tip-Tip distance and the distance from the residues in the luminal region. For the subsequent OPES simulations, 16 configurations were randomly selected from the REST2 ensemble, equilibrated, and used as initial states. The same initial configurations were used for the WT simulations after mutating I392E and S426A back.

The OPES runs employed 16 walkers, with biases imposed on the collective variables of a single protomer and the K^+^ ion. The barrier parameter was set to 12 kcal mol^-1^, and hills were deposited every 2 ps. Hill information was shared among replicas every 10 ps. Each OPES walker was simulated for 200 ns. A spring constant of 20 kcal mol^-1^ Å^-2^ was used for the cylindrical restraint of K^+^.

### Ancestral Protein Reconstruction

To build a seed alignment for sequence retrieval and alignment, PROMALS3D^101^ was used to generate a structure-based alignment (default settings, MAFFT aligned^102^) using representatives of the SLC1 family: hEAAT1, hEAAT2, hEAAT3, hASCT1, dEAAT2, and Glt_Ph_. Tails were manually trimmed from the resulting alignment and sequences were retrieved from the Reference Proteomes^103^ database using jackHMMER^104^ with the seed alignment. The search converged after 5 iterations and yielded 10,395 sequences from the sodium:dicarboxylate symporter family; hits were annotated with taxonomic information using E-Utilities^105^ and filtered using seqkit^106^ to remove incomplete/ambiguous sequences (‘X’ residues, < 70% query coverage). Sequences were clustered to 80% identity and 80% length using mmseqs^107^, resulting in 1,086 clusters. Sequences of interest (hEAAT1-5, dEAAT1-2) were used as representatives for their respective clusters.

The cluster representatives were aligned with HMMER^108^ using a profile built from the seed alignment, and insertions different from the seed profile were removed. The resulting alignment was manually curated to remove additional truncated/gappy sequences and extended N- and C-termini, resulting in 631 sequences for subsequent tree analysis. A preliminary tree was run using VeryFastTree^109^ to assess overall topology; long branches and underrepresented phyla (*Porifera* = 1; *Orthonectida* = 1; *Placozoa* = 2) were flagged using Pykit^110^ and removed. Subsequent maximum-likelihood trees were run using IQ-TREE^111–113^ with a *Cnidarian* outgroup; once the optimal model (LG) was identified using ModelFinder, long branches and rogue taxa were iteratively removed using Phykit, RogueNaRok^114^, and manually. The final tree was determined from 100 total tree runs, with the number of rate categories empirically determined for each run using a range of 6-14; the best tree used for subsequent analysis used an LG+F+R12 model. Finally, bootstrap values were determined using UFBoot2, and ancestral states were determined using the IQ-TREE implementation.

## Acknowledgements.

This research was supported by the Australian Government through an Australian Research Council Future Fellowship FT220100717 (R.M.R) and a National Health and Medical Research Council Project Grant APP1164494 (R.M.R). This study was supported by the National Institutes of Health (NIH) through grants R24-GM145965 (E.T.), R01-DK135088 (E.T.) and R35-GM165890 (K.D.R.). We are grateful to the laboratory animal services of the Victor Chang Cardiac Research Institute for the maintenance of the *Xenopus laevis* facility and supply of oocytes. We acknowledge the use of instruments at the Monash Ramaciotti Centre for Cryo-Electron Microscopy, a Node of Microscopy Australia and the Monash University MASSIVE high-performance computing facility for data management. The authors acknowledge the technical and scientific assistance of Sydney Microscopy & Microanalysis, the University of Sydney node of Microscopy Australia. Computational simulations were conducted using resources provided by the National Science Foundation Supercomputing Centers (ACCESS grant number MCA06N060) and the Texas Advanced Computing Center (TACC) at The University of Texas at Austin (TACC grant number MCB24004). Bioinformatics analyses were performed on the Central Instructional and Research Computing Environment (CIRCE) cluster computer at the University of South Florida. This research was conducted by the Australian Research Council Industrial Transformation Training Centre for Cryo-Electron Microscopy of Membrane Proteins for Drug Discovery (IC200100052).

## Author contributions

Q.W. and R.M.R. conceived the study. Q.W., C.B., and R.M.R. designed and performed functional experiments in oocytes. Y.J., H.V., Q.W., and R.M.R. designed and performed biochemistry and cryo-EM experiments and analysis. A.F., S.P., and E.T. designed and performed simulation experiments. M.D. and K.D.R. designed and performed bioinformatic analyses. The manuscript was written by Q.W., Y.J., C.B. and R.M.R. with input from all authors.

## Competing interests

The authors declare no competing interests.

## Materials & Correspondence

Renae M Ryan

## Notes

### Competing Interest Statement

The authors have declared no competing interest.

### Summary of Updates

The order of authors was corrected.

