## Extended Data for "One high-order ion-coupling mechanism generates multiple SLC1A stoichiometries"

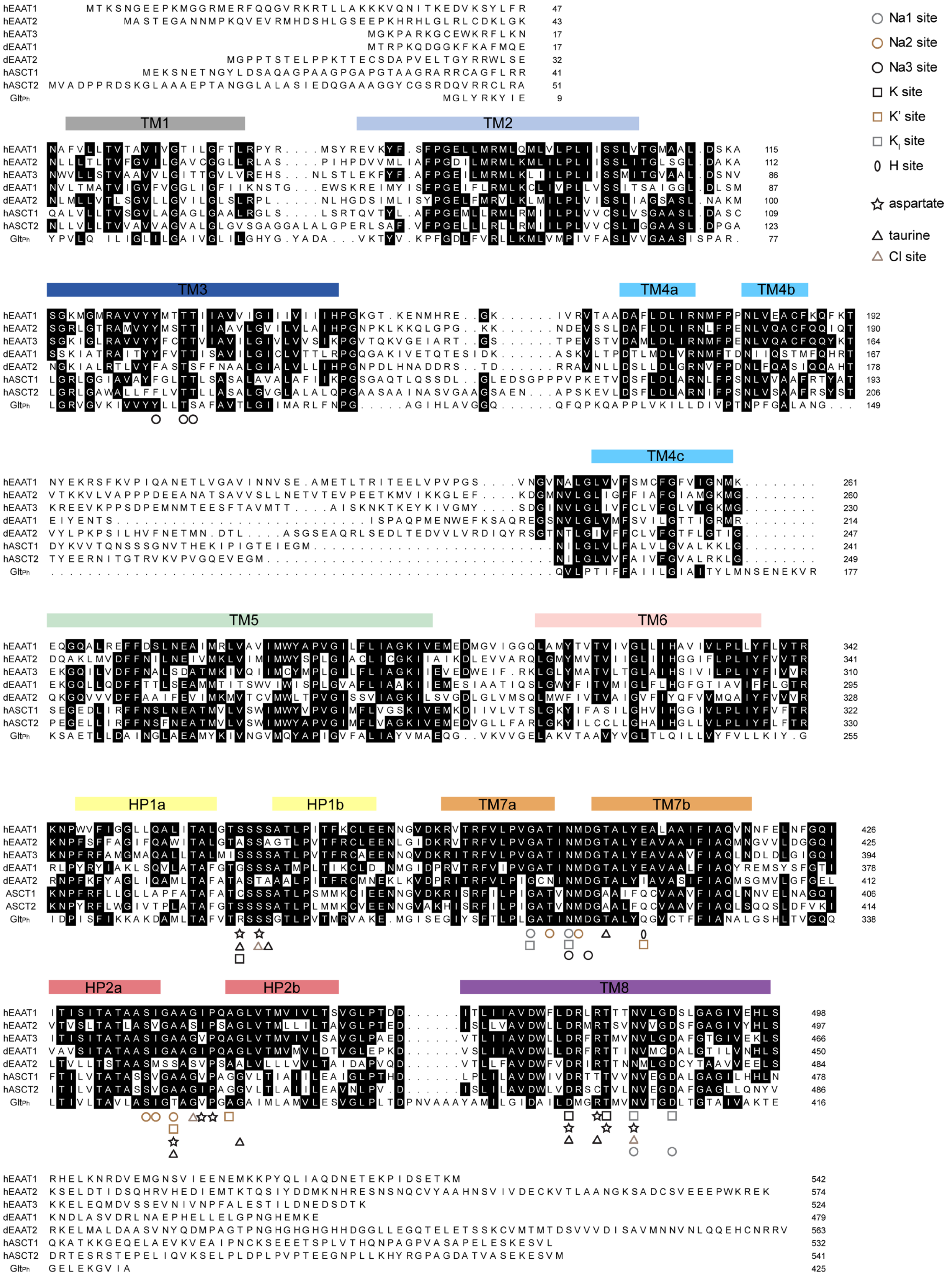


**Extended Data Figure 1 | Sequence alignment of SLC1A family members from different organisms.** Sequence identification code (UniProt): hEAAT1 (P43003-1), hEAAT2 (P43004-1), hEAAT3 (P43005-1), dEAAT1 (O77062-1), dEAAT2 (Q9VPS6-1), hASCT1 (P43007-1), hASCT2 (Q15758-1) and Glt$_{Ph}$ (O59010-1) and binding sites indicated.

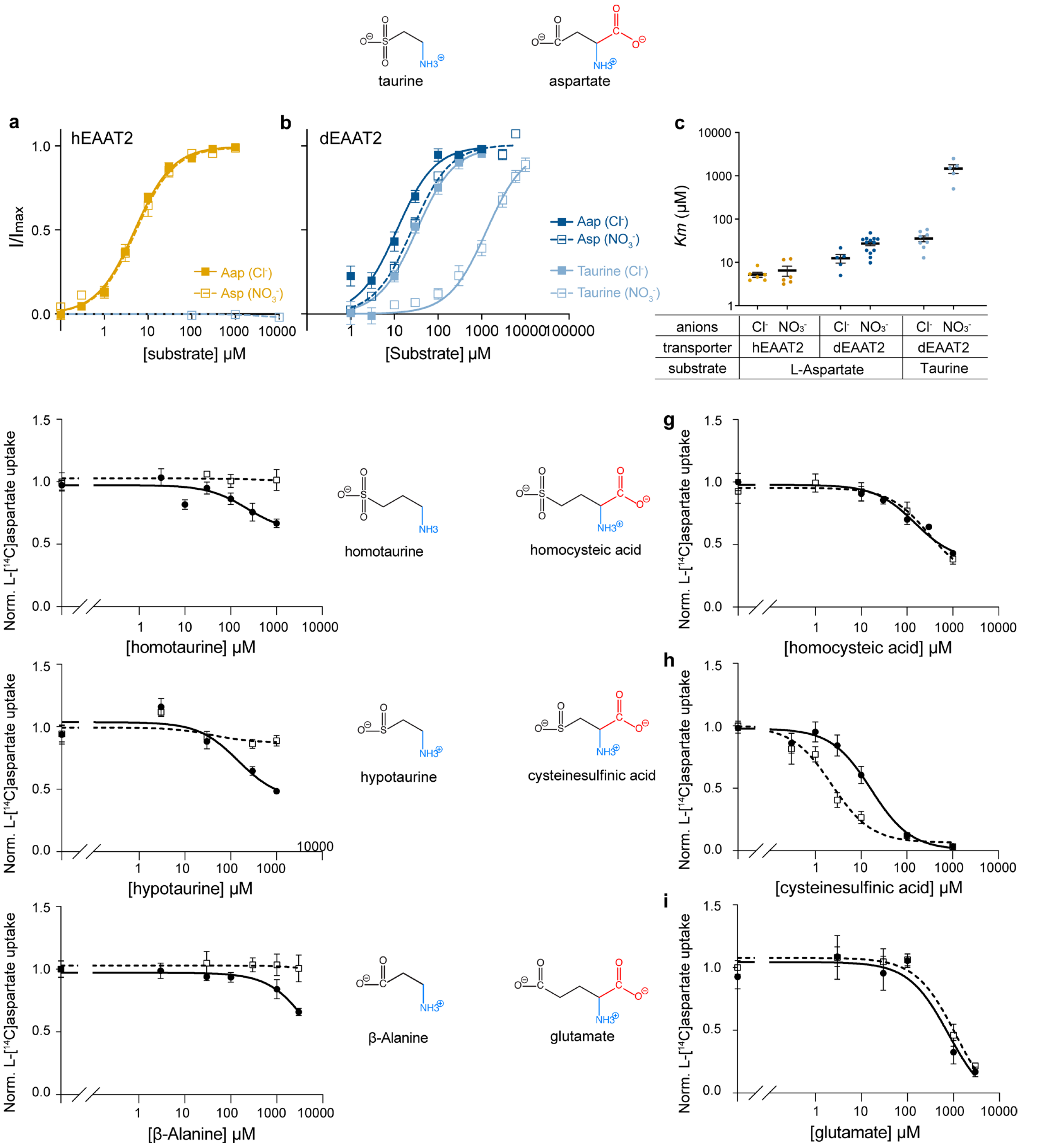


**Extended Data Figure 2 | Cl⁻ dependence of dEAAT2 is determined by the presence or absence of the α-carboxyl group.** Aspartate and taurine titration via hEAAT2 (**a**) and dEAAT2 (**b**) were conducted in chloride buffer (filled squares) or nitrate buffer (open squares). The corresponding apparent affinities of each condition ($K_M$) are plotted (**c**). Inhibition of L-[14C]aspartate uptake by taurine (**d-f**) or aspartate derivatives (**g-i**) with or without a second carboxyl group on the α-carbon (highlighted in red) were tested in the absence (open symbols) or presence (closed symbols) of $Cl^-$. Methanesulfonate ($MES^-$) was used as a $Cl^-$ substituent to maintain the equivalent molarity in the $Cl^-$ free buffer. Data presented as mean ± SEM ) and analysed using GraphPad Prism 10. Measurements were obtained from two different batches of oocytes. Precise $IC_{50}$ values provided in **Extended Data Table 2** and exact n values provided in **Supplementary Table 2.**

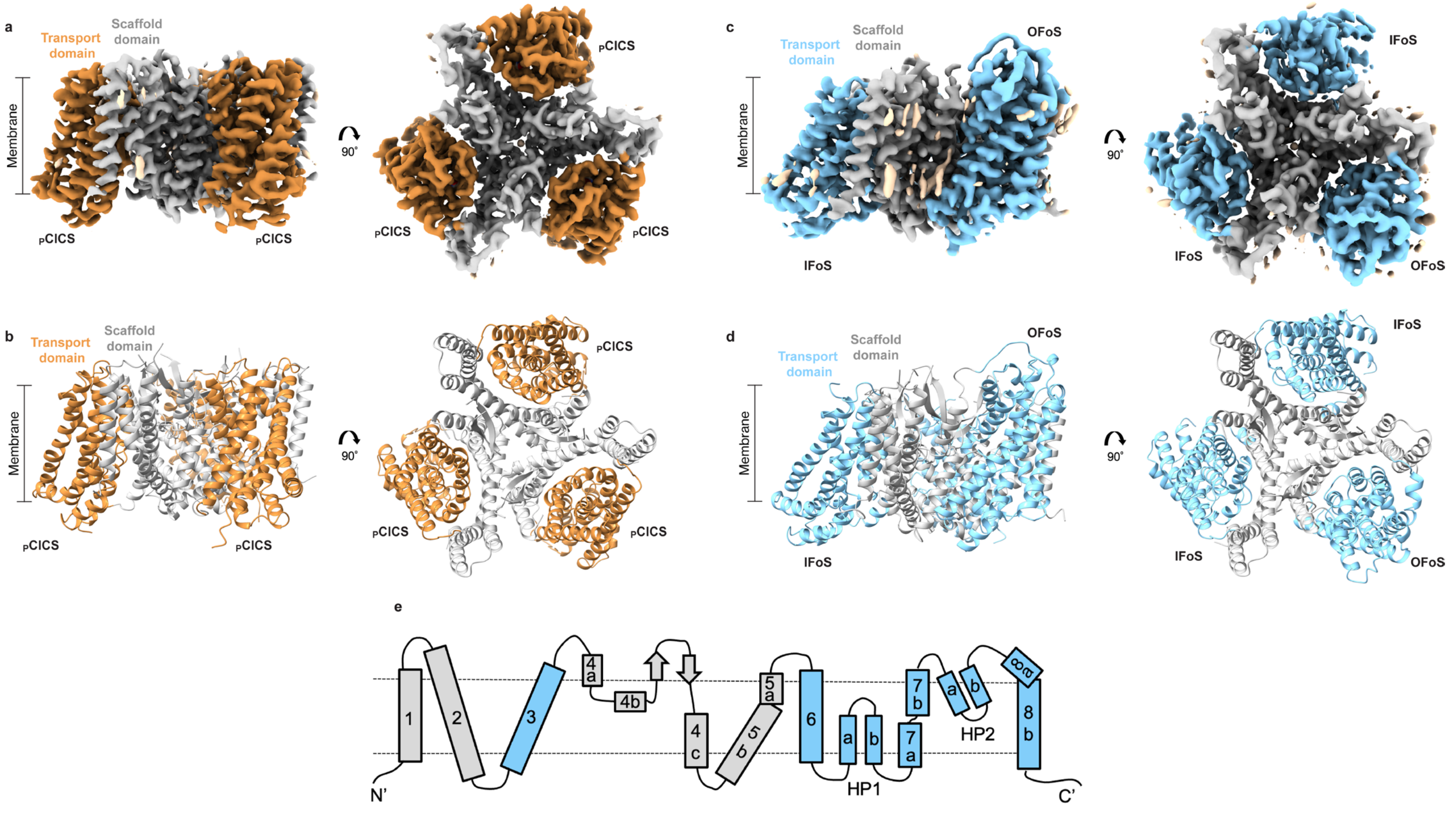


**Extended Data Figure 3 | Cryo-EM structure of dEAAT2.** (**a**) Cryo-EM density map of dEAAT2 bound to taurine at 2.4 Å resolution, shown in side (left) and top-down (right) view. Each protomer is coloured, transport domain (orange) and scaffold domain (grey). All three protomers adopt the pre-open chloride-conducting state conformation ($_{P}$ClCS). (**b**) Side view (left) and top-down (right) view of the atomic model of the taurine-bound dEAAT2 trimer, with domains coloured as in (**a**). (**c**) 2.64 Å cryo-EM density map of dEAAT2 trimer shown in side (left) and top-down (right) view. Each protomer is coloured, transport domain (cornflower blue) and scaffold domain (grey). One protomer adopts the outward-facing open state (OF$_{O}$S), while the other two are in the inward-facing open state (IF$_{O}$S). **d** Atomic model of dEAAT2 trimer in side (left) and top-down (right) view, with domains coloured as in (**c**). (**e**) Schematic topology of the secondary structure of a dEAAT2 protomer. Transmembrane helices are labelled 1–8, with domains colored as in (**c**).

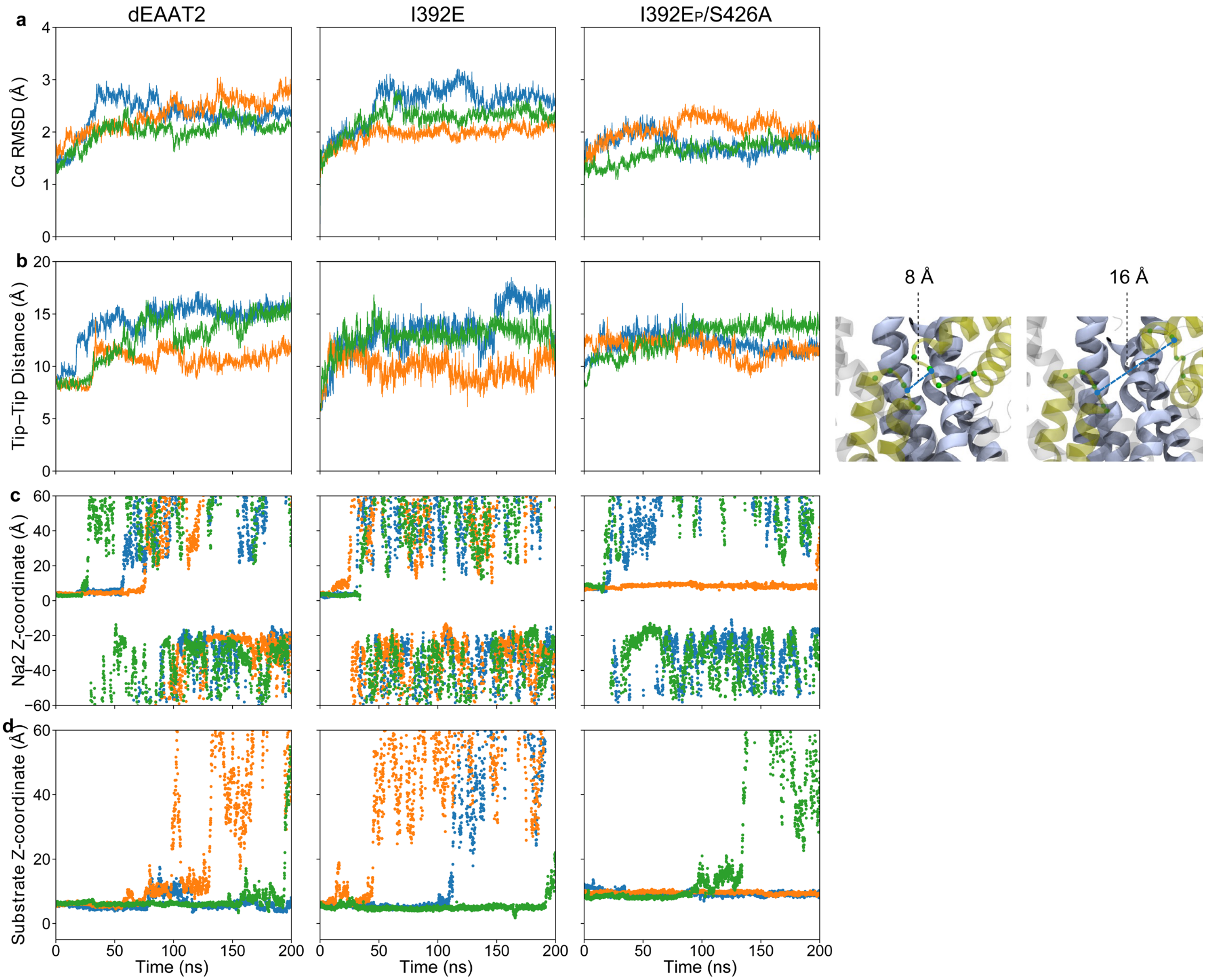


**Extended Data Figure 4 | Equilibrium structural dynamics of the OF$_C$S.** Shown are 200 ns unconstrained equilibrium MD simulations for three variants: dEAAT2, I392E$_{dE2}$ (unprotonated), and I392E$_P$/S426A (protonated). Each column corresponds to one variant, and each row represents a measured quantity. (**a**) Cα RMSD of each system compared to the OF$_C$S structure. (**b**) Tip–tip distance, defined as the distance between the mass centers (schematically indicated as blue circles) of the Cα atoms of residues 348–352 (HP1) and 426–430 (HP2). Green, orange, and blue traces represent protomers A, B, and C, respectively. Sample representation of the OF$_C$S I392E-S426A with (left) tip-tip distance at 8 Å and (right) at 16 Å. The scaffold domain, transport domain, HP1, and HP2 are shown in gray, cornflower blue, and gold, respectively. The Cα atoms are shown as green spheres. (**c**) z-coordinate of $Na^+$ at the Na2 site. (**d**) Z-coordinate of the substrate (aspartate). Green, orange, and blue colors represent $Na^+$ and substrate in protomers A, B, and C, respectively.

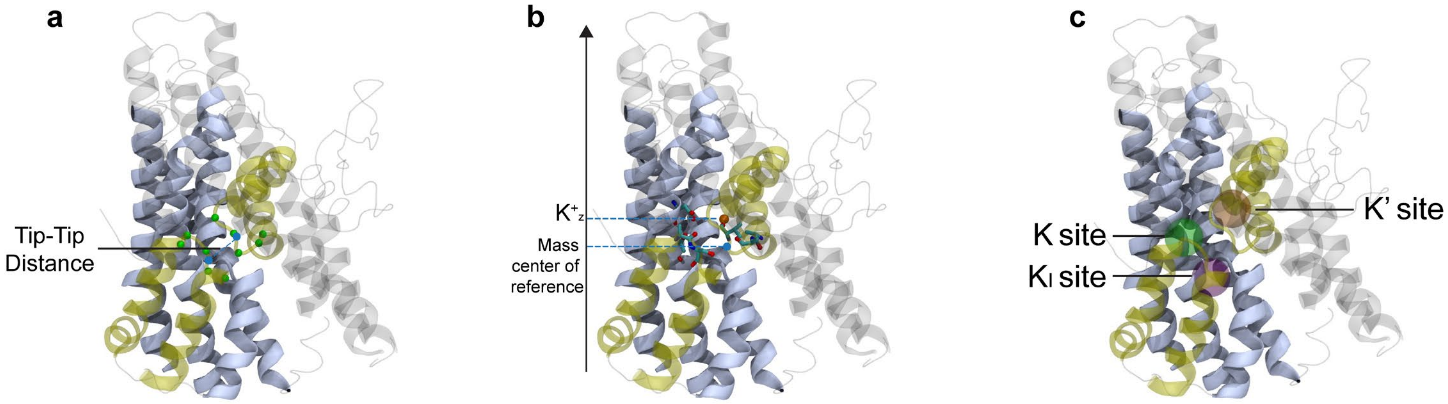


**Extended Data Figure 5 | Representation of collective variables used in the biased MD study.**
(**a**) Representation of the $_P$ClCS and schematic illustration of the tip–tip distance. The tip–tip distance is defined as the distance between the mass centers of the Cα atoms of residues 348–352 (HP1) and 426–430 (HP2). The mass centers are schematically indicated by blue circles, and the corresponding Cα atoms are shown as green spheres. (**b**) The z-distance is defined as the distance along the membrane normal (z-axis) between the $K^+$ ion and the mass center of the side-chain oxygen atoms of residues D462 and T466 (TM8), A349 and S350 (HP1), and A426, A427, and V429 (HP2). The $K^+$ ion is shown as an orange sphere, the selected residues are depicted in licorice representation, and the oxygen mass center is schematically indicated by a blue circle. (**c**) Schematic representation of the identified potential $K^+$ binding sites, $K_I$, K, and K'. The scaffold domain, transport domain, HP1, and HP2 are shown in gray, cornflower blue, and gold, respectively.

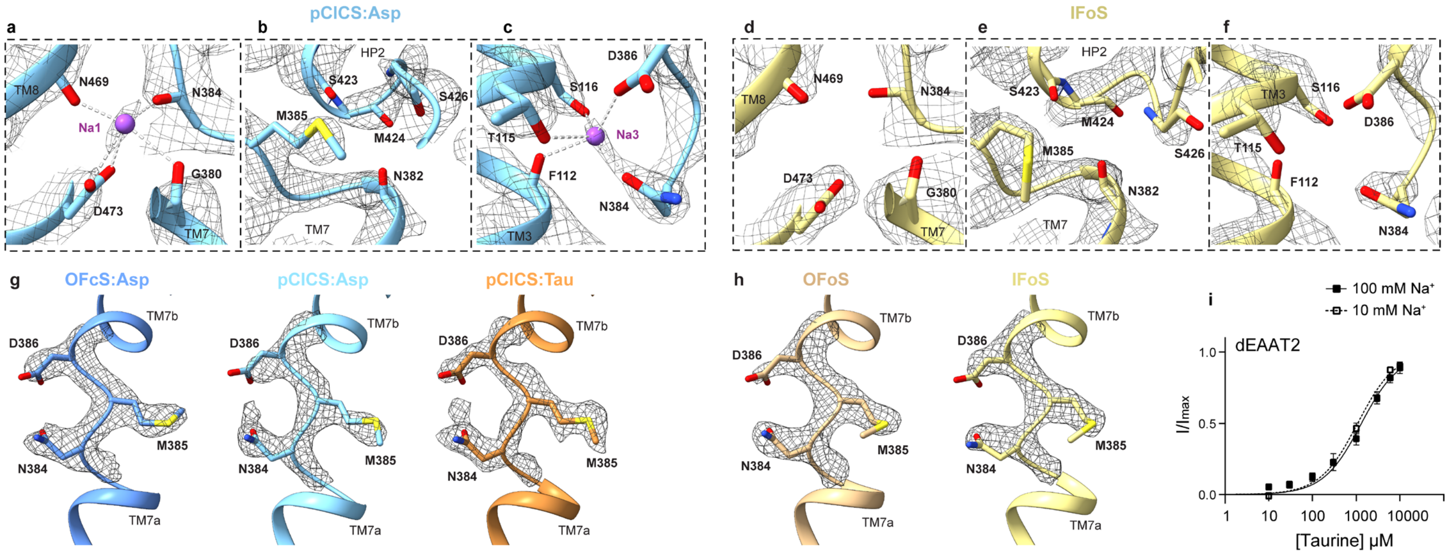


**Extended Data Figure 6 | Cryo-EM structures show coordination of $Na^+$ binding sites.** Na1 (left panel), Na2 (middle panel) and Na3 (right panel). $_{P}$ClCS-Asp (a-c), and $IF_OS$ (d-f). Position of M385 from the highly conserved NMD motif is similar in both closed (g) and open states (h). Taurine dose-response curves for dEAAT2 in the presence of 10 mM $Na^+$ (open squares) and 100 mM $Na^+$ (filled squares) (i).

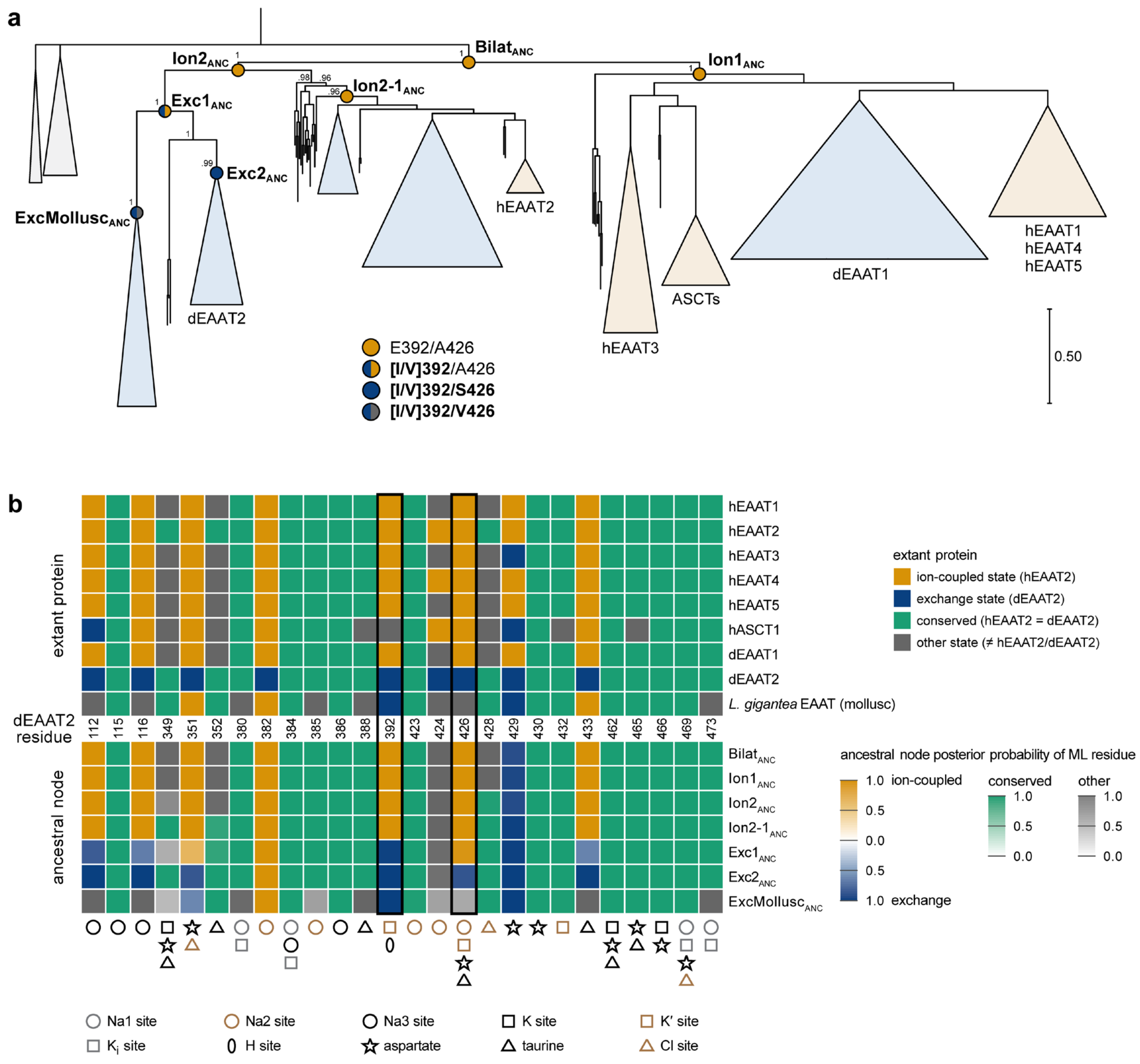


**Extended Data Figure 7. A protostomic lineage of ion-independent exchangers that emerged from ion-coupled transporters.** (a) Reduced maximum-likelihood (ML) phylogeny of SLC1 transporters in metazoans, rooted to cnidarians. Numbers near key branches represent bootstrap support values. The collapsed clades are proportional to the number of comprising sequences and are colored by taxonomy (skyblue, protostomes; sand, deuterostomes). The left and right halves of each node represent the ancestral states at positions $392_{dE2}$ and $426_{dE2}$, respectively. (b) Key sequence transitions from ion-coupled to ion-independent transporters, in both extant and ancestral SLCs. hEAAT2 is the reference for the ion-coupled state, and dEAAT2 is the reference for the exchange state. Top panel displays extant SLC1 transporters containing either hEAAT2-specific residues (yellow), dEAAT2-specific residues (blue), conserved residues (green), or other residues (grey). The bottom panel shows if ML residues in the displayed ancestral sequences match hEAAT2 and/or dEAAT2, along with the corresponding posterior probabilities.

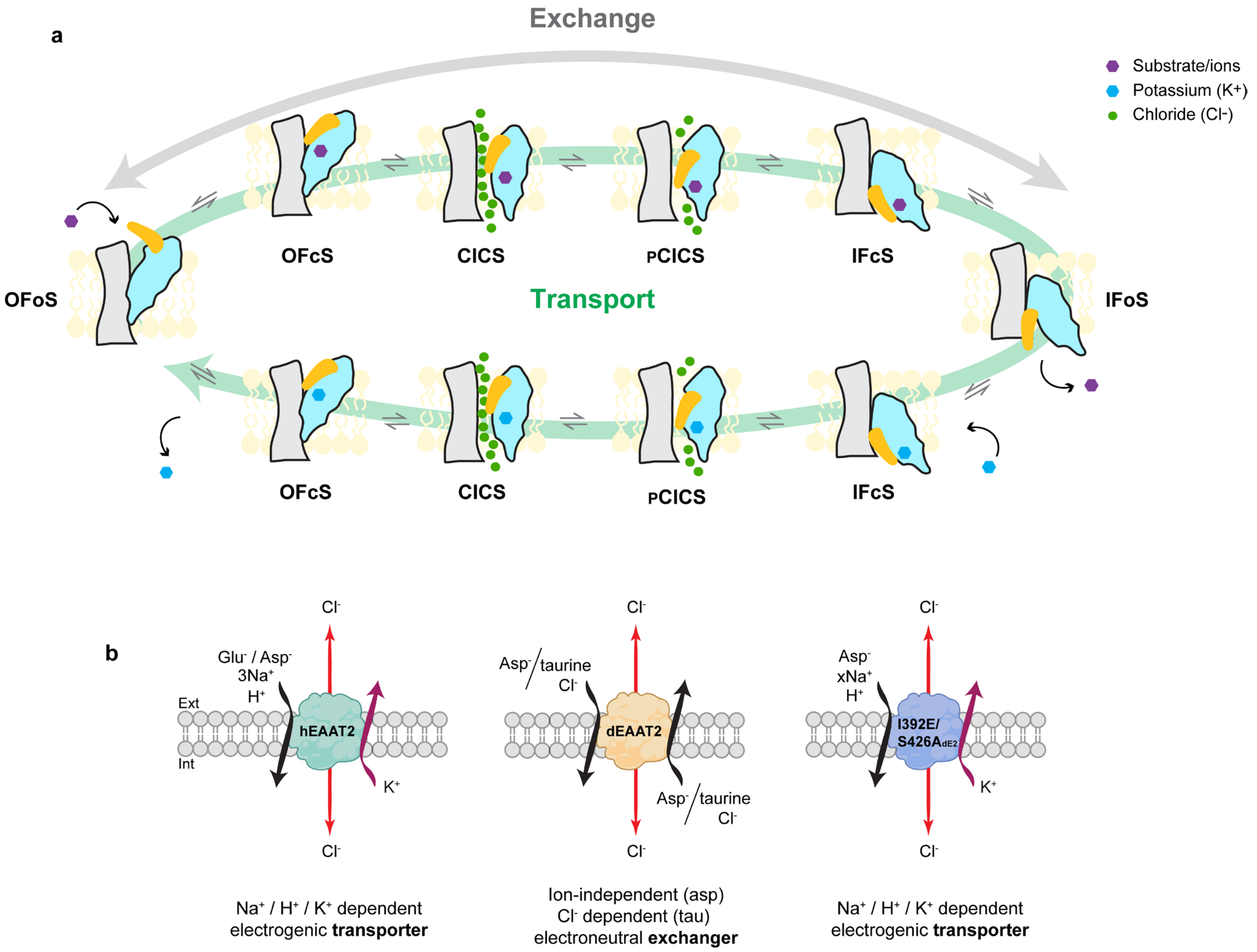


**Extended Data Figure 8 | Transport cycle of hEAAT, dEAAT2 and I392E/S426A$_{dE2}$. (a)** All three transporters can undergo exchange (grey arrow) if substrate is available on both sides of the membrane. The transporter moves from OF$_{O}$S to the IF$_{O}$S through a series of intermediate states including the $_{P}$ClCS described in this study. For exchange, hEAAT2 requires $Na^+$ and substrate, while dEAAT2 can undergo exchange in the absence of $Na^+$. Only hEAAT2 and I392E/S426A$_{dE2}$ can undergo the complete transport cycle (green arrow) where $K^+$ binding is required for the transporter to return to the OF$_{O}$S. (**b**) The stoichiometry of transport via hEAAT2, exchange via dEAAT2 and transport via I392E/S426A$_{dE2}$.

**Extended Data Table 1 | Properties of hEAAT2 and dEAAT2 measured in buffers containing various anions and $Na^+$ concentrations.** Currents were fitted to Michaelis-Menten by least squares from which apparent affinities ($K_m$) and maximal currents ($I_{max}$) were obtained. Data was collected from two different batches of oocytes and presented as mean ± SEM, with the number of cells (n) used indicated.

| Transporter | Substrate | $Na^+$ (mM) | Anion | $K_m$ (μM) | $I_{max}$ (nA) | n |
|---|---|---|---|---|---|---|
| **hEAAT2** | L-Aspartate | 100 | $Cl^-$ | 5.2 ± 0.7 | 76.6 ± 13.4 | 6 |
| | | | $Br^-$ | 9.4 ± 1.2 | 189.2 ± 31.6 | 7 |
| | | | $I^-$ | 5.3 ± 0.6 | 216.2 ± 20.3 | 7 |
| | | | $NO_3^-$ | 6.5 ± 1.7 | 610.0 ± 90.9 | 6 |
| | | 10 | $NO_3^-$ | 422.8 ± 56.5 | 381.4 ± 57.7 | 5 |
| **dEAAT2** | L-Aspartate | 100 | $Cl^-$ | 12.4 ± 2.8 | 32.2 ± 4.1 | 5 |
| | | | $Br^-$ | 12.7 ± 3.2 | 47.1 ± 8.7 | 5 |
| | | | $I^-$ | 25.7 ± 5.2 | 510.1 ± 62.1 | 5 |
| | | | $NO_3^-$ | 27.3 ± 3.1 | 826.1 ± 128.0 | 13 |
| | | 10 | $NO_3^-$ | 20.2 ± 1.1 | 634.1 ± 84.6 | 10 |
| | Taurine | 100 | $Cl^-$ | 29.2 ± 4.6 | 31.0 ± 9.1 | 6 |
| | | | $Br^-$ | 20.9 ± 3.3 | 32.8 ± 3.4 | 5 |
| | | | $I^-$ | 80.3 ± 14.1 | 176.8 ± 16.5 | 7 |
| | | | $NO_3^-$ | 1464.8 ± 328.1 | 851.7 ± 128.1 | 5 |
| | | 10 | $NO_3^-$ | 1099.5 ± 135.8 | 533.1 ± 84.2 | 5 |
| **I392E/S426A$_{dE2}$** | L-Aspartate | 100 | $NO_3^-$ | 31.7 ± 12.7 | 137 ± 38.8 | 5 |
| | | 10 | | 231.7 ± 59.8 | 185 ± 60.1 | 6 |
| **I392E$_{dE2}$** | L-Aspartate | 100 | $NO_3^-$ | 12.3 ± 2.7 | 161.8 ± 9.7 | 5 |
| | | 10 | | 19.0 ± 4.9 | 187.9 ± 23.9 | 5 |
| **S426A$_{dE2}$** | L-Aspartate | 100 | $NO_3^-$ | 20.6 ± 8.6 | 149.9 ± 24.3 | 5 |
| | | 10 | | 20.9 ± 4.6 | 287.6 ± 54.6 | 5 |

**Extended Data Table 2 | $IC_{50}$ (μM) of compounds that inhibit L-[[14]C]aspartate uptake via dEAAT2.** Analysed data from **Extended Data Fig. 2** assayed in the absence or presence of $Cl^-$. 95% confidence intervals are shown in brackets. Exact number of oocytes tested are indicated Supplementary Table 2. NA (not applicable).

| Taurine derivatives | | | Aspartate derivatives | | |
|---|---|---|---|---|---|
| | | $IC_{50}$ (μM) | | | $IC_{50}$ (μM) |
| taurine | *Cl⁻* | 32.3 (18.2 – 59.9) | cysteic acid | *Cl⁻* | 1.0 (0.7 to 1.3) |
| | *Cl⁻ free* | NA | | *Cl⁻ free* | 1.0 (0.7 – 1.6) |
| homotaurine | *Cl⁻* | 230.0 (38.7 – 3301.0) | homocysteic acid | *Cl⁻* | 147.6 (49.0 – 571.4) |
| | *Cl⁻ free* | NA | | *Cl⁻ free* | 299.0 (67.6 – ∞) |
| hypotaurine | *Cl⁻* | 146.0 (33.9 – 905.6) | cysteinesulfinic acid | *Cl⁻* | 16.4 (9.8 – 29.1) |
| | *Cl⁻ free* | NA | | *Cl⁻ free* | 2.3 (1.5 – 3.4) |
| β-alanine | *Cl⁻* | 5253 (190.5 – ∞) | glutamate | *Cl⁻* | 780.2 (271.9 – 2724) |
| | *Cl⁻ free* | NA | | *Cl⁻ free* | 1129 (279.4 – 8440) |
