## Supplementary Data for "One high-order ion-coupling mechanism generates multiple SLC1A stoichiometries"

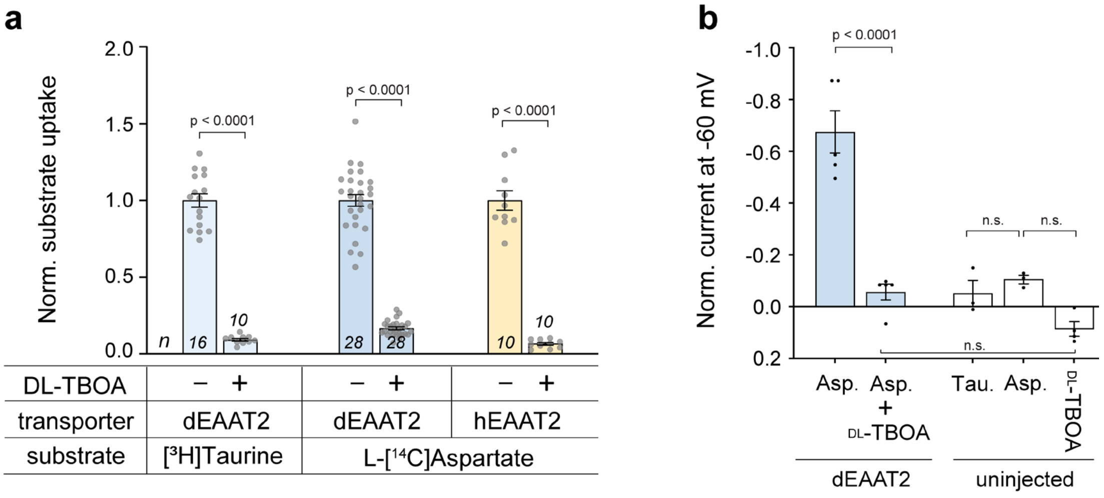

**Supplementary Figure 1** (**a**) The pan-EAAT inhibitor DL-TBOA (100 μM) blocks radiolabelled L-[$^{14}$C]aspartate and [$^{3}$H]taurine transport via dEAAT2 and L-[$^{14}$C]aspartate transport via hEAAT2 (10 μM). (**b**) Application of DL-TBOA (30 μM) inhibits aspartate (15 μM)-activated currents via dEAAT2. 1 mM taurine, aspartate, and 30 μM DL-TBOA alone did not elicit significant currents in uninjected cells. Currents normalized to the maximum current elicited by 15 μM aspartate via dEAAT2. The exact number of oocytes used (n), from 2-3 different batches, is indicated as dots.

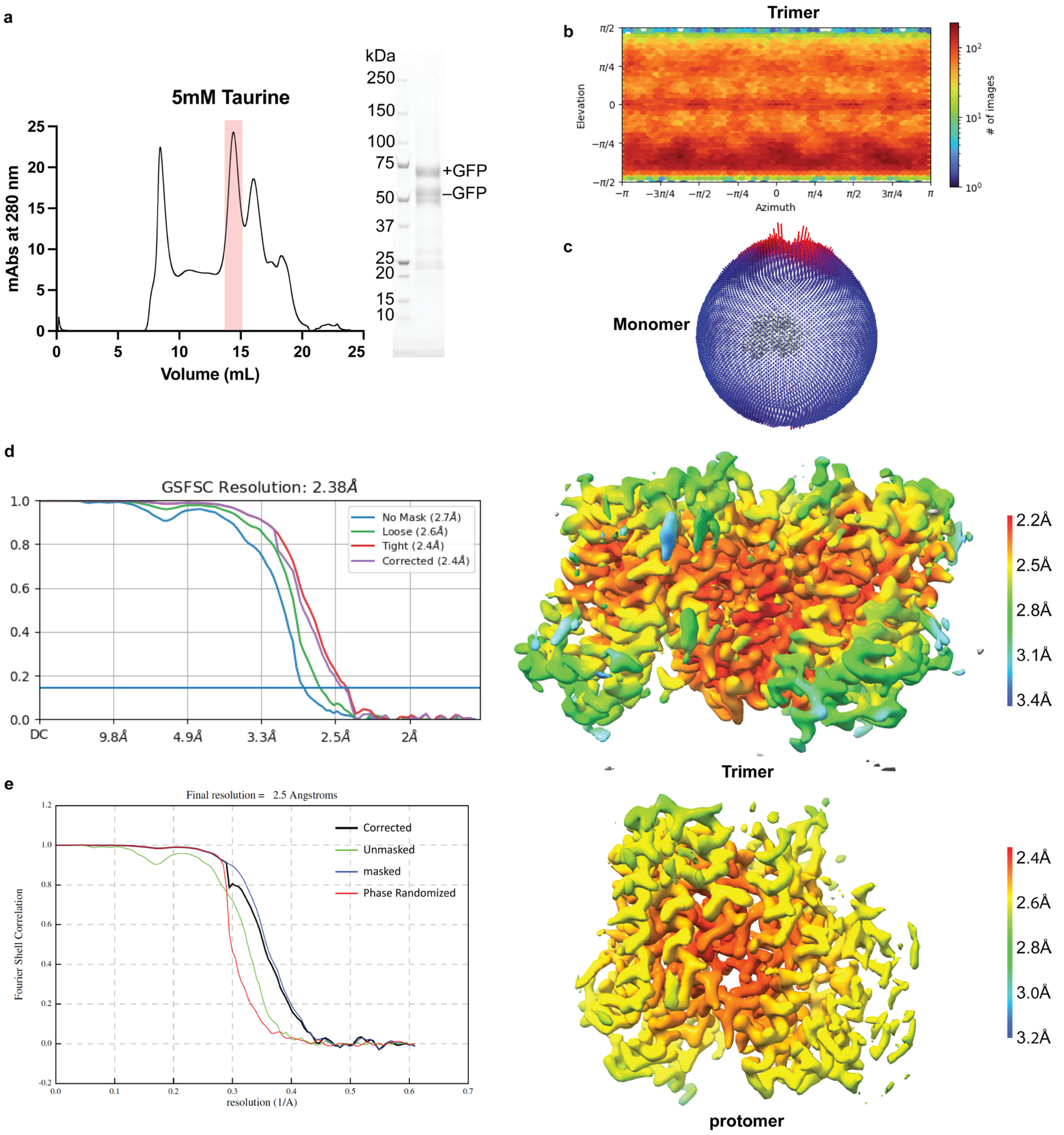

**Supplementary Figure 2 | Cryo-EM analysis of dEAAT2 in the presence of 5 mM taurine. (a)** Size-exclusion chromatograms of dEAAT2 in 5 mM taurine. The shaded peak indicates the fraction collected for cryo-EM studies. The inset shows the SDS-PAGE analysis of the concentrated peak fractions, revealing monomeric dEAAT2 protein with and without a C-terminal GFP tag. The bands corresponding to dEAAT2 are clearly observed between 50 and 75 kDa. **(b)** Angular distribution heatmaps calculated using cryoSPARC for trimeric dEAAT2 particles. The heatmaps display the particle distribution across viewing angles, with higher particle counts indicated by warmer colors. **c** Particle orientation plots from the 3D reconstructions of inward-facing occluded monomer. Fourier Shell Correlation (FSC) curves (left) and local resolution maps (right) for the 3D reconstructions of the inward-facing trimeric (**d**) and inward-facing monomeric (**e**) conformations of dEAAT2. The FSC plots provide an estimate of the resolution, and the local resolution maps highlight regional variations in reconstruction quality.

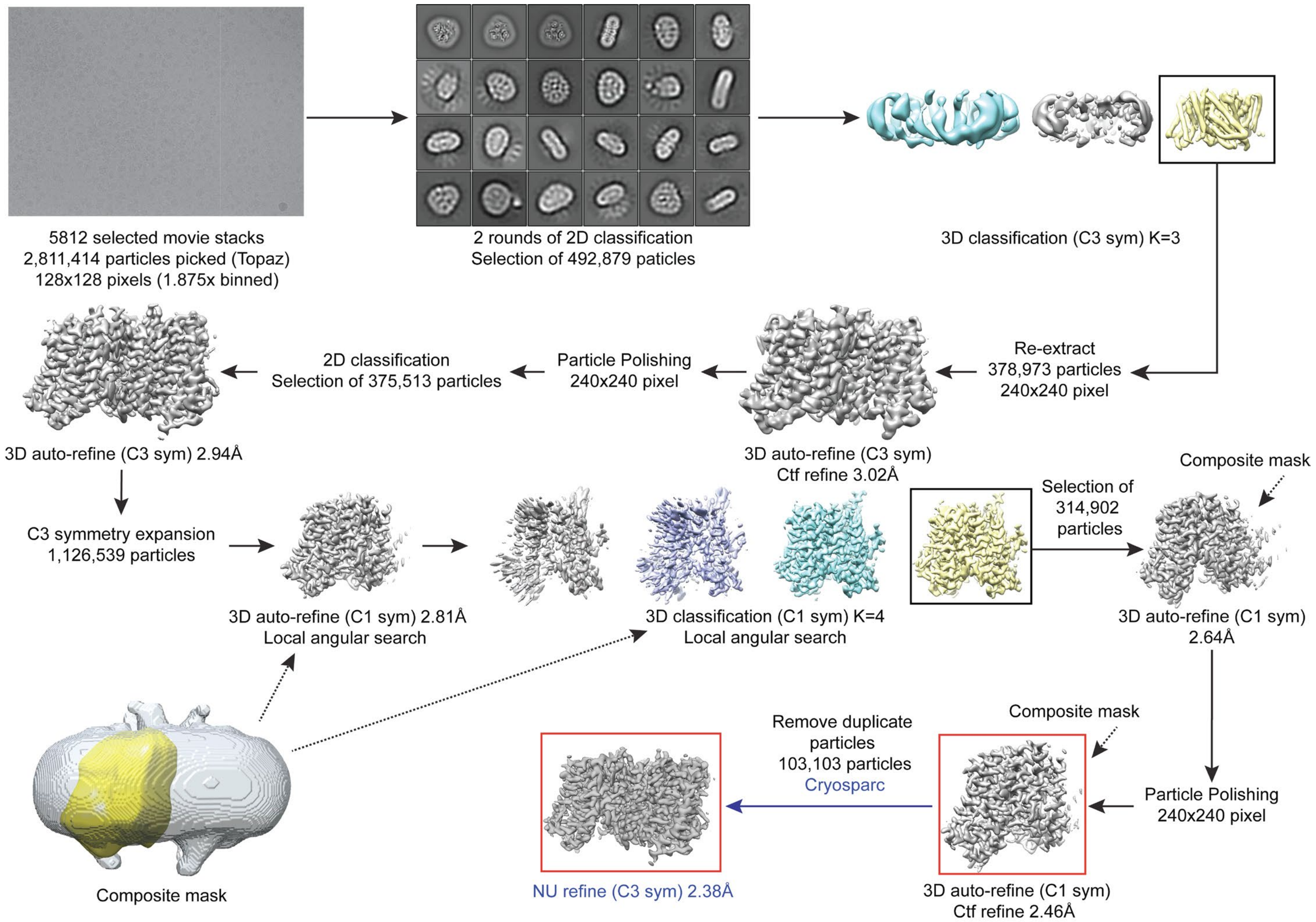

**Supplementary Figure 3 | Cryo-EM data processing workflow for dEAAT2 in 5 mM taurine.** The figure outlines the data processing pipeline for 3D reconstruction of dEAAT2 in the presence of 5 mM taurine. The 3D reconstructions are presented without B-factor sharpening. Reconstructions highlighted in red boxes were selected for atomic model building.

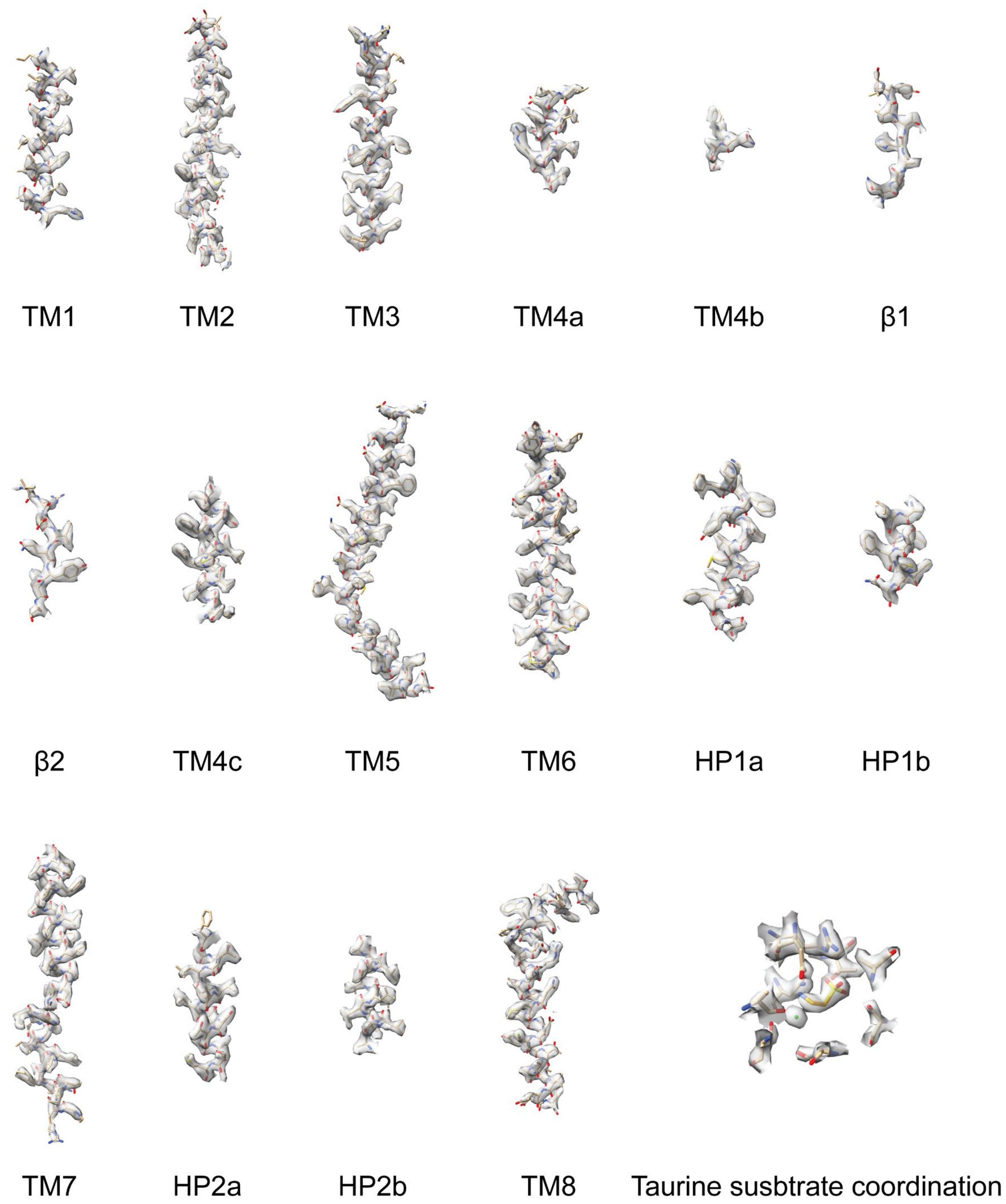

**Supplementary Figure 4 | Cryo-EM maps for representative regions of dEAAT2 with taurine**. The cryo-EM maps for the TM1-TM8 helices, the HP1, the HP2 and the substrate coordiantion site.

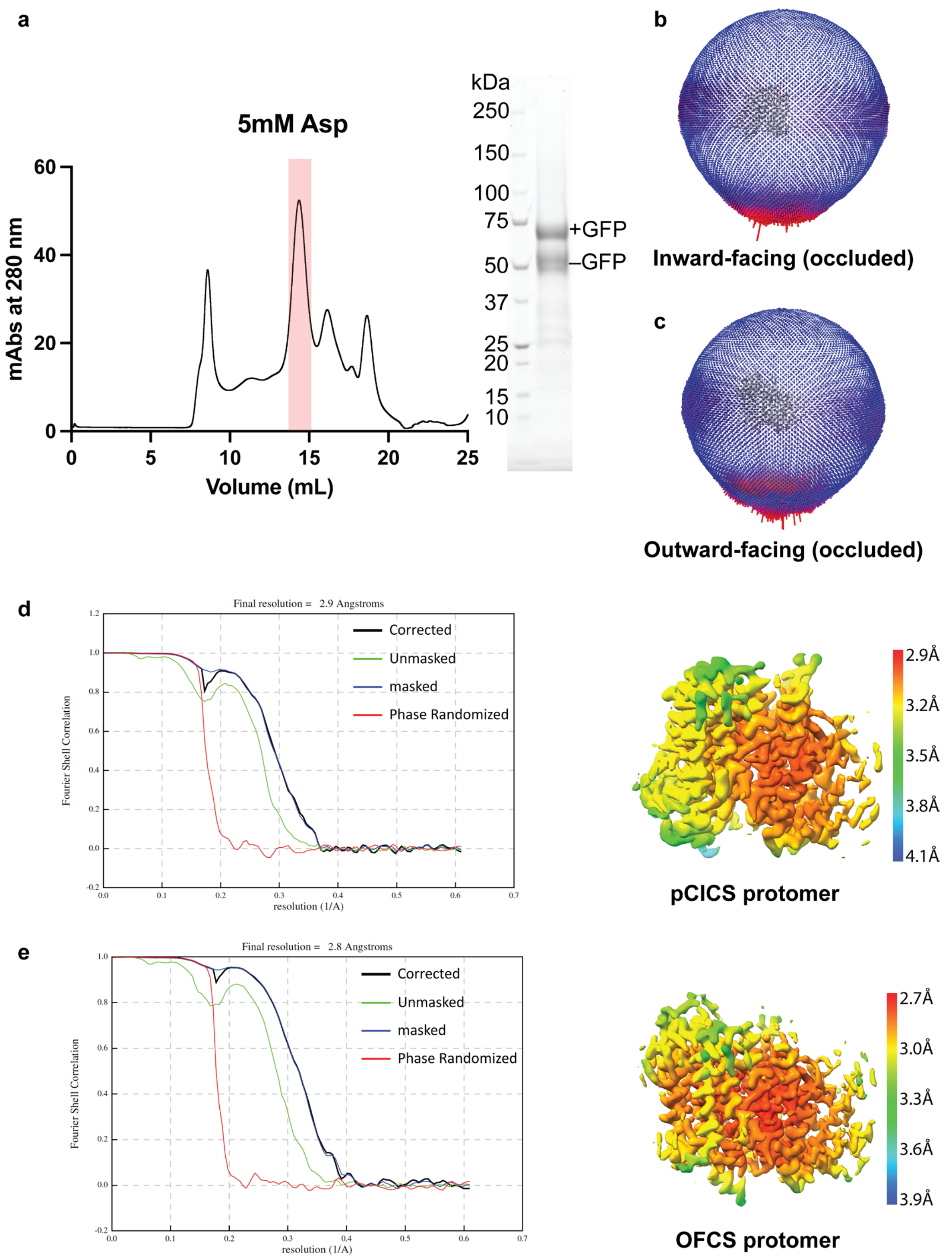

**Supplementary Figure 5 | Cryo-EM analysis of dEAAT2 in the presence of 5 mM aspartate. (a)** Size-exclusion chromatogram of dEAAT2 in 5 mM aspartate. The shaded region indicates the fraction collected for cryo-EM analysis. The inset shows SDS-PAGE of the concentrated peak fractions, revealing monomeric dEAAT2 protein with and without a C-terminal GFP tag. dEAAT2 bands are observed between 50 and 75 kDa. Particle orientation plots from the 3D reconstructions of inward-facing occluded monomer (**b**) and outward-facing occluded monomer (**c**) conformations. Fourier Shell Correlation (FSC) curves (left) and local resolution maps (right) for the inward-facing occluded monomer (**d**) and outward-facing occluded monomer (**e**) conformations of dEAAT2. FSC plots estimate the global resolution, and local resolution maps highlight regional variations in reconstruction quality.

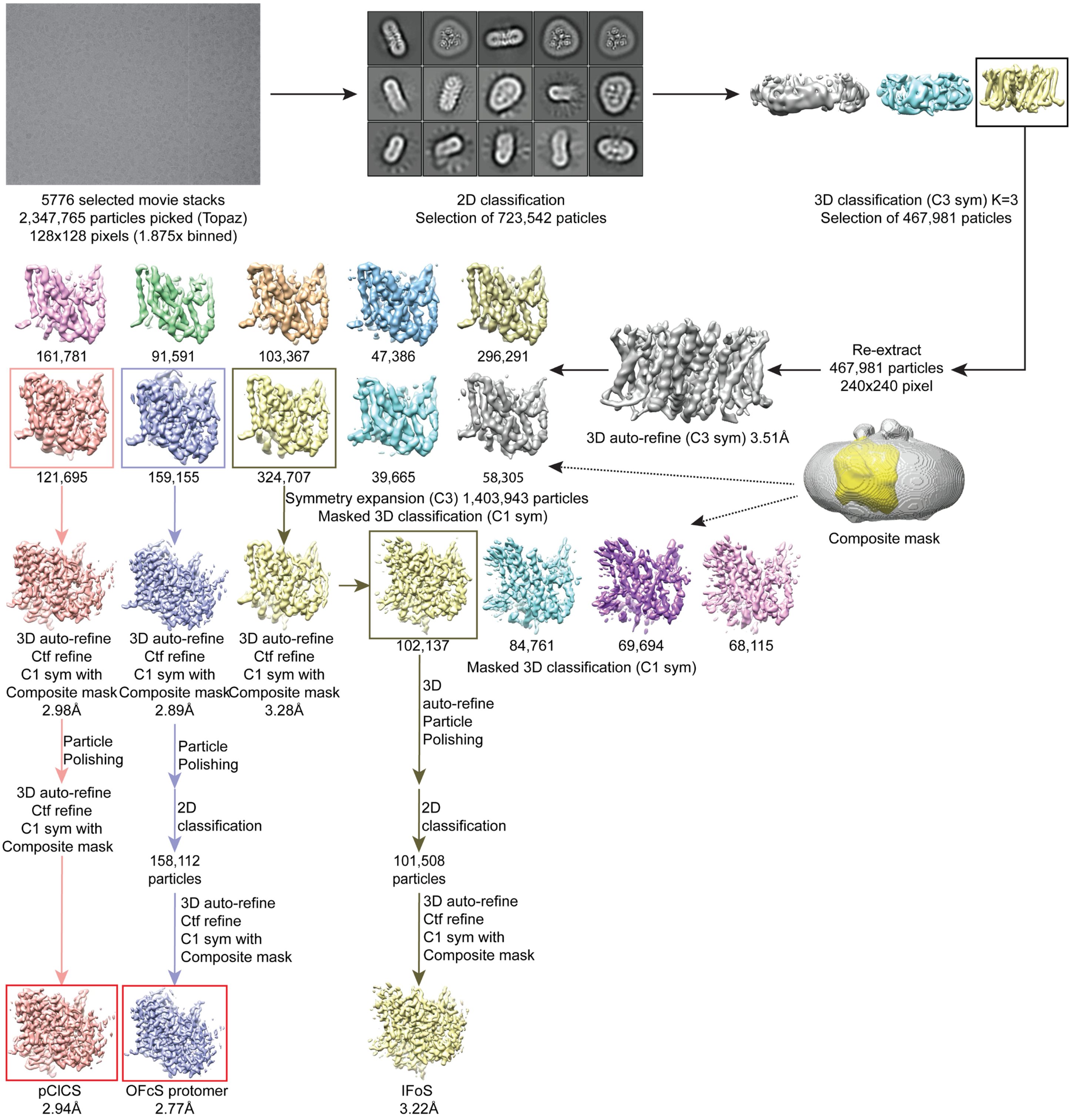

**Supplementary Figure 6 | Cryo-EM data processing workflow for dEAAT2 in 5 mM aspartate.** The figure outlines the data processing pipeline for 3D reconstruction of dEAAT2 in the presence of 5 mM aspartate. The 3D reconstructions are presented without B-factor sharpening. Reconstructions highlighted in red boxes were selected for atomic model building.

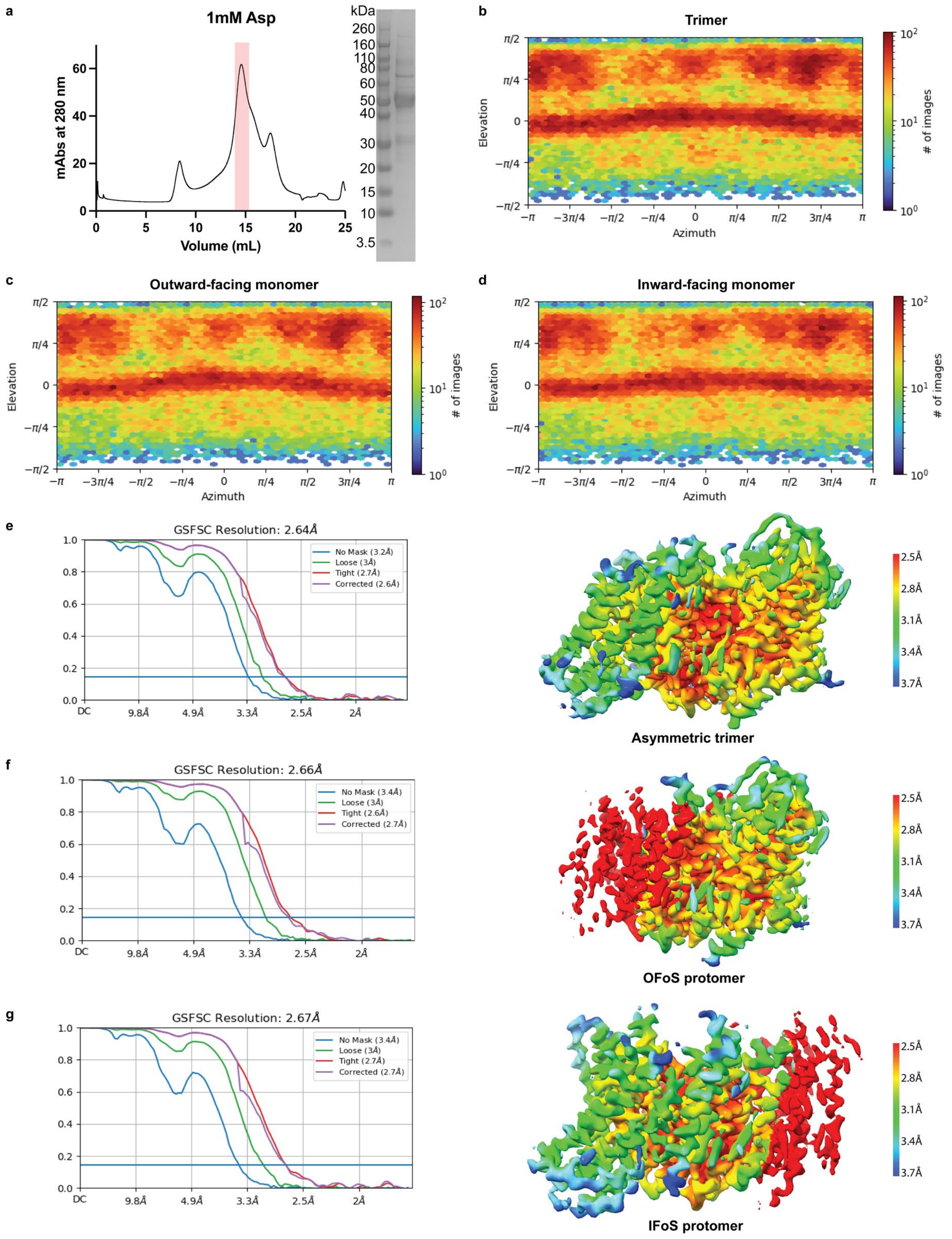

**Supplementary Figure 7 | Cryo-EM analysis of dEAAT2 in the presence of 1 mM aspartate. (a)** Size-exclusion chromatogram of dEAAT2 in 1 mM aspartate. The shaded region indicates the fraction collected for cryo-EM analysis. The inset shows SDS-PAGE of the concentrated peak fractions, revealing monomeric dEAAT2 protein with bands between 50 and 75 kDa. Angular distribution heatmaps calculated in cryoSPARC for trimeric (**b**), outward-facing monomeric (**c**), and inward-facing monomeric (**d**) dEAAT2 particles. Warmer colors indicate higher particle counts. Fourier Shell Correlation (FSC) curves (left) and local resolution maps (right) for the 3D reconstructions of hybrid outward-facing/inward-facing trimeric (**e**), outward-facing open monomeric (**f**), and inward-facing open monomeric (**g**) dEAAT2 conformations. FSC plots indicate global resolution, while local resolution maps highlight regional reconstruction quality.

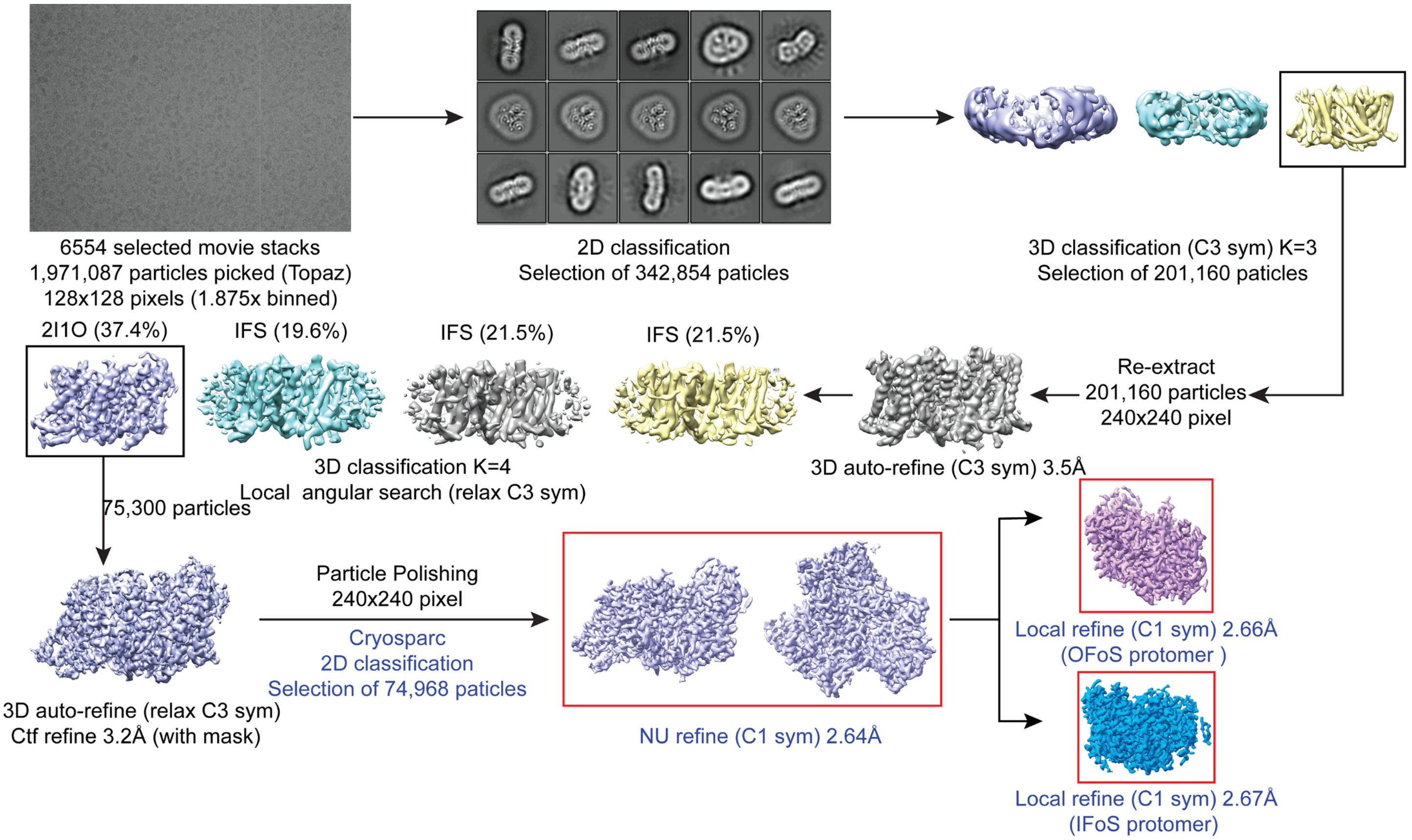

**Supplementary Figure 8 | Cryo-EM data processing workflow for dEAAT2 in 1 mM aspartate.** The figure outlines the data processing pipeline for 3D reconstruction of dEAAT2 in the presence of 1 mM aspartate. The 3D reconstructions are presented without B-factor sharpening. Reconstructions highlighted in red boxes were selected for atomic model building.

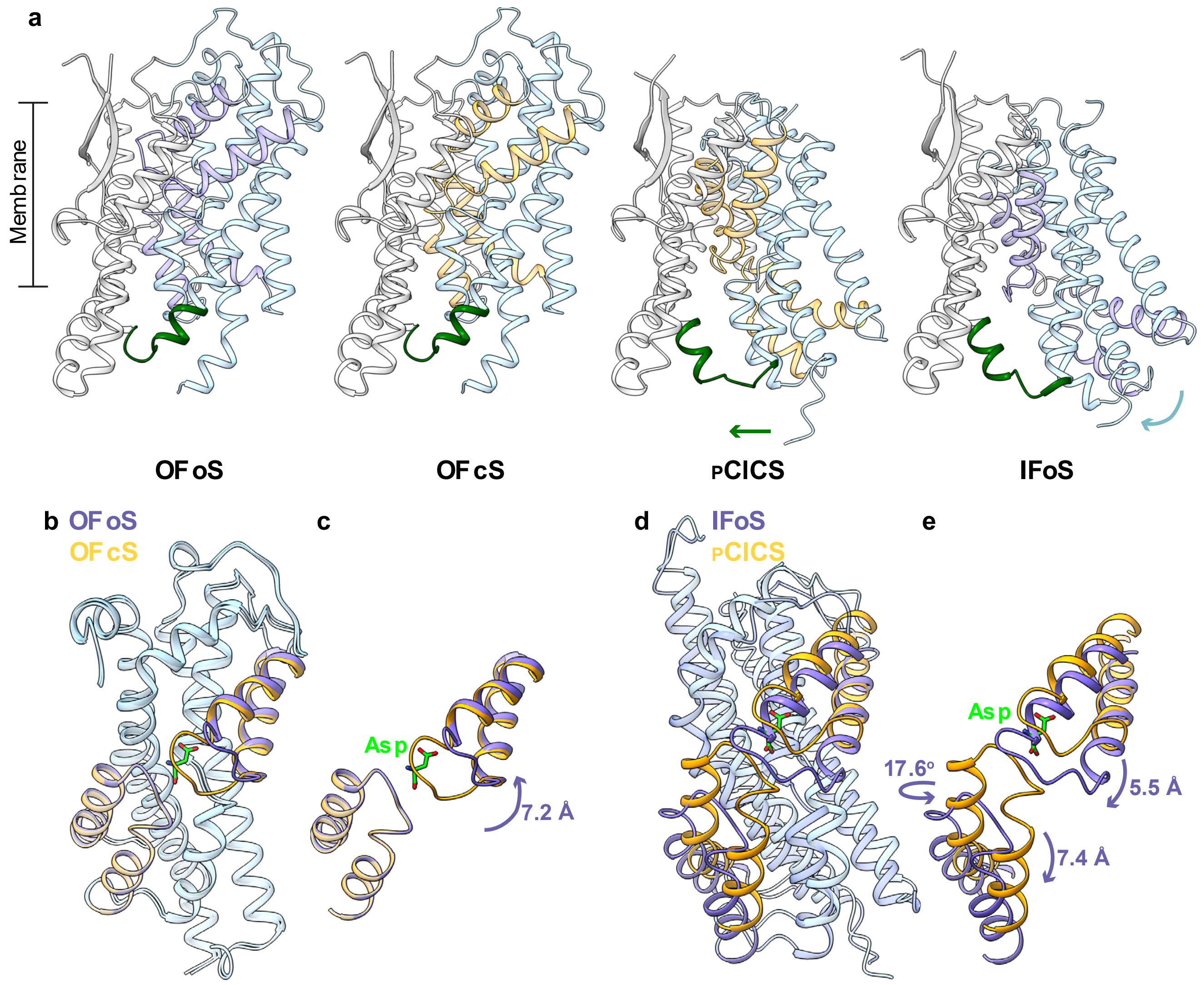

**Supplementary Figure 9 | Structural comparison of dEAAT2 in outward-facing and inward-facing states, illustrating the elevator transport mechanism.** (**a**) Structural alignment of the transport domain in four conformational states: outward-facing open ($OF_OS$), outward-facing closed ($OF_CS$), pre-Chloride Conducting State ($_P$CLCS), and inward-facing open ($IF_OS$). Residues 93–103, which transition from TM3 in the inward-facing state to TM2 in the outward-facing state, are highlighted in dark green. The transport domain is shown in cornflower blue, the scaffold domain in grey, HP1 and HP2 are gold (closed states) and light purple (open states). (**b**) Superposition of the transport domain of dEAAT2 $OF_OS$ and $OF_CS$, demonstrating the 7.2 Å opening of HP2. (**d**) Superposition of the transport domain of dEAAT2 $_P$CLCS and $IF_OS$, demonstrating the downward movement of HP1 (7.4 Å) and HP2 (5.5 Å) and a 17.6º rotation of the transport domain. (**c-e**) TMD's 3,5,7,8 removed for clarity. (**b-e**) The transport domain is shown in cornflower blue, HP1 and HP2 are gold (closed states) and light purple (open states), substrate aspartate is shown as green sticks (**b-e**).

**Supplementary Figure 10 | Convergence of constant-pH MD and OPES free-energy calculations. (a,b)** Running pKa estimates as a function of cumulative constant-pH MD simulation time for the investigated protonatable residues in **(a)** dEAAT2 and **(b)** I392E/S426A. Equivalent sites in the three protomers were treated as replicate simulations and combined for pKa estimation, as described in Methods. The bulk aspartate reference used for calibration is also shown. Shaded bands indicate the standard deviation across the three protomer replicas. Unshown curves correspond to pKa estimates outside the sampled pH range of the constant-pH calculations and were not interpreted quantitatively. **(c)** Self-convergence of the two-dimensional OPES free-energy landscapes for dEAAT2 and I392E/S426A. Self-convergence was quantified using the RMSD of the free-energy landscape reconstructed at cumulative simulation time relative to the final free-energy landscape. The landscapes were evaluated on the same discretized two-dimensional collective-variable space and vertically aligned before calculating the RMSD to account for the arbitrary additive constant in free energy. Simulation time represents cumulative sampling across the 16 OPES walkers. The dashed line indicates $k_B T$ at 310 K. The RMSD falls below this threshold after approximately 2.0 μs for dEAAT2 and 2.2 μs for I392E/S426A and remains below it for the remainder of the simulations.

**Supplementary Table 1. Cryo-EM data collection, refinement and validation statistics.**

| | dEAAT2 pClCS trimer in complex with taurine (EMD-73466) (PDB 9YTL) | dEAAT2 pClCS protomer in complex with taurine (EMD-73467) (PDB 9YTM) |
|---|---|---|
| **Data collection and processing** | | |
| Magnification | 105,000× | |
| Voltage (kV) | 300 | |
| Electron exposure (e–/Å$^2$) | 60 | |
| Defocus range (μm) | –0.4 to –1.4 | |
| Pixel size (Å) | 0.82 | |
| Symmetry imposed | C3 | C1 |
| Initial particle images (no.) | 2,811,414 | 2,811,414 |
| Final particle images (no.) | 211,799 | 314,902 |
| Map resolution (Å) | 2.38 | 2.46 |
| FSC threshold | 0.143 | 0.143 |
| Map resolution range (Å) | 2.1 – 7.4 | 2.44 – 3.36 |
| **Refinement** | | |
| Initial model used (PDB code) | dEAAT2 pClCS protomer | model built de novo from cryo-EM density |
| Model resolution (Å) | 2.5 | 2.6 |
| FSC threshold | 0.5 | 0.5 |
| Map sharpening *B* factor (Å$^2$) | –86.8 | –76.7 |
| Model composition | | |
| Nonhydrogen atoms | 9,528 | 3,176 |
| Protein residues | 1,248 | 416 |
| Ligands | TAU:3/NA:9/CL:3 | TAU:1/NA:3/CL:1 |
| *B* factors (Å$^2$) | | |
| Protein (mean) | 49.71 | 49.78 |
| Ligand (mean) | 34.85 | 36.29 |
| R.m.s. deviations | | |
| Bond lengths (Å) | 0.012 | 0.012 |
| Bond angles (°) | 1.785 | 1.786 |
| Validation | | |
| MolProbity score | 0.83 | 0.87 |
| Clashscore | 1.18 | 1.38 |
| Poor rotamers (%) | 0.00 | 0.00 |
| Ramachandran plot | | |
| Favored (%) | 98.54 | 98.29 |
| Allowed (%) | 1.46 | 1.71 |
| Disallowed (%) | 0.00 | 0.00 |

**Supplementary Table 1. Cryo-EM data collection, refinement and validation statistics (continued)**

| | dEAAT2 $_{P}$ClCS protomer in complex with aspartate (EMD-73464) (PDB 9YTJ) | dEAAT2 OF$_{C}$S protomer in complex with aspartate (EMD-73465) (PDB 9YTK) |
|---|---|---|
| **Data collection and processing** | | |
| Magnification | 105,000× | |
| Voltage (kV) | 300 | |
| Electron exposure (e–/Å$^2$) | 60 | |
| Defocus range (μm) | –0.4 to –1.4 | |
| Pixel size (Å) | 0.82 | |
| Symmetry imposed | C1 | C1 |
| Initial particle images (no.) | 2,347,765 | 2,347,765 |
| Final particle images (no.) | 159,155 | 121,695 |
| Map resolution (Å) | 2.94 | 2.77 |
| FSC threshold | 0.143 | 0.143 |
| Map resolution range (Å) | 2.98–6.46 | 2.74–6.35 |
| **Refinement** | | |
| Initial model used (PDB code) | model built de novo from cryo-EM density | model built de novo from cryo-EM density |
| Model resolution (Å) | 3.1 | 2.9 |
| FSC threshold | 0.5 | 0.5 |
| Map sharpening *B* factor (Å$^2$) | –87.4 | –76.8 |
| Model composition | | |
| Nonhydrogen Atoms | 3113 | 3302 |
| Protein residues | 408 | 431 |
| Ligands | ASP:1/NA:2 | ASP:1/NA:3 |
| *B* factors (Å$^2$) | | |
| Protein (mean) | 59.71 | 35.30 |
| Ligand (mean) | 20.00 | 20.00 |
| R.m.s. deviations | | |
| Bond lengths (Å) | 0.012 | 0.012 |
| Bond angles (°) | 1.807 | 1.807 |
| Validation | | |
| MolProbity score | 0.78 | 0.70 |
| Clashscore | 0.94 | 0.59 |
| Poor rotamers (%) | 0.87 | 0.27 |
| Ramachandran plot | | |
| Favored (%) | 98.51 | 98.36 |
| Allowed (%) | 1.49 | 1.64 |
| Disallowed (%) | 0.00 | 0.00 |

**Supplementary Table 1. Cryo-EM data collection, refinement and validation statistics (continued)**

| | dEAAT2 asymmetric trimer (EMD-73461) (PDB 9YTG) | dEAAT2 $OF_OS$ protomer (EMD-73462) (PDB 9YTH) | dEAAT2 $IF_OS$ protomer (EMD-73463) (PDB 9YTI) |
|---|---|---|---|
| **Data collection and processing** | | | |
| Magnification | | 105,000× | |
| Voltage (kV) | | 300 | |
| Electron exposure (e–/Å$^2$) | | 60 | |
| Defocus range (μm) | | –0.4 to –1.4 | |
| Pixel size (Å) | | 0.82 | |
| Symmetry imposed | C1 | C1 | C1 |
| Initial particle images (no.) | 2,811,414 | 2,811,414 | 2,811,414 |
| Final particle images (no.) | 74,968 | 74,968 | 74,968 |
| Map resolution (Å) | 2.64 | 2.66 | 2.67 |
| FSC threshold | 0.143 | 0.143 | 0.143 |
| Map resolution range (Å) | 2.4 – 8 | 2.1 – 4.1 | 2.5 – 5.5 |
| **Refinement** | | | |
| Initial model used (PDB code) | dEAAT2 OFoS and IFoS structures | model built de novo from cryo-EM density | model built de novo from cryo-EM density |
| Model resolution (Å) | 2.9 | 2.9 | 3.0 |
| FSC threshold | 0.5 | 0.5 | 0.5 |
| Map sharpening *B* factor (Å$^2$) | –73.3 | –83.7 | –81.9 |
| Model composition | | | |
| Nonhydrogen atoms | 9584 | 3302 | 3141 |
| Protein residues | 1,259 | 433 | 413 |
| Ligands | NA:2 | NA:2 | 0 |
| *B* factors (Å$^2$) | | | |
| Protein (mean) | 102.09 | 99.98 | 92.67 |
| Ligand (mean) | 112.44 | 112.44 | – |
| R.m.s. deviations | | | |
| Bond lengths (Å) | 0.012 | 0.011 | 0.012 |
| Bond angles (°) | 1.790 | 1.761 | 1.800 |
| Validation | | | |
| MolProbity score | 0.97 | 0.86 | 0.76 |
| Clashscore | 2.03 | 1.33 | 0.46 |
| Poor rotamers (%) | 0.66 | 0.55 | 0.57 |
| Ramachandran plot | | | |
| Favored (%) | 98.15 | 98.37 | 97.54 |
| Allowed (%) | 1.85 | 1.63 | 2.46 |
| Disallowed (%) | 0.00 | 0.00 | 0.00 |

**Supplementary Table 2 | Number of repeats (n) for dose-response curves constructed from radiolabelled uptake experiments** from **Extended Data Fig. 2** and **Fig. 2e-g.** Exact number of oocytes tests (n) are reported for each compound or $Cl^-$ concentration. Data was collected from two different batches of oocytes.

| | | | | | | | | | | |
|---|---|---|---|---|---|---|---|---|---|---|
| | | [compounds] µM, n | | | | | | | | |
| taurine | $Cl^-$ | 0 µM, 23 | 1 µM, 9 | 10 µM, 9 | 30 µM, 9 | 100 µM, 9 | 300 µM, 21 | 1000 µM, 6 | | |
| | $Cl^-$ free | 0 µM, 28 | 30 µM, 6 | 300 µM, 12 | 1000 µM, 8 | | | | | |
| homotaurine | $Cl^-$ | 0 µM, 31 | 3 µM, 14 | 10 µM, 9 | 30 µM, 16 | 100 µM, 17 | 300 µM, 17 | 1000 µM, 17 | | |
| | $Cl^-$ free | 0 µM, 6 | 30 µM, 8 | 100 µM, 8 | 1000 µM, 8 | | | | | |
| hypotaurine | $Cl^-$ | 0 µM, 8 | 3 µM, 8 | 30 µM, 8 | 300 µM, 8 | 1000 µM, 8 | | | | |
| | $Cl^-$ free | 0 µM, 15 | 3 µM, 7 | 30 µM, 8 | 300 µM, 8 | 1000 µM, 16 | | | | |
| β-alanine | $Cl^-$ | 0 µM, 7 | 3 µM, 6 | 30 µM, 7 | 100 µM, 7 | 1000 µM, 7 | 3000 µM, 7 | | | |
| | $Cl^-$ free | 0 µM, 7 | 30 µM, 7 | 300 µM, 7 | 1000 µM, 7 | 3000 µM, 7 | | | | |
| cysteic acid | $Cl^-$ | 0 µM, 27 | 0.3 µM, 8 | 1 µM, 17 | 3 µM, 8 | 10 µM, 17 | 30 µM, 9 | 100 µM, 16 | 300 µM, 17 | 1000 µM, 18 |
| | $Cl^-$ free | 0 µM, 18 | 0.3 µM, 8 | 1 µM, 8 | 3 µM, 8 | 10 µM, 7 | 100 µM, 8 | 1000 µM, 10 | | |
| homocysteic acid | $Cl^-$ | 0 µM, 19 | 10 µM, 9 | 30 µM, 9 | 100 µM, 9 | 300 µM, 9 | 1000 µM, 18 | | | |
| | $Cl^-$ free | 0 µM, 7 | 1 µM, 6 | 10 µM, 8 | 100 µM, 7 | 1000 µM, 7 | | | | |
| cysteinesulfinic acid | $Cl^-$ | 0 µM, 27 | 0.3 µM, 7 | 1 µM, 16 | 3 µM, 16 | 10 µM, 16 | 100 µM, 15 | 1000 µM, 17 | | |
| | $Cl^-$ free | 0 µM, 27 | 0.3 µM, 8 | 1 µM, 16 | 3 µM, 15 | 10 µM, 13 | 100 µM, 16 | 1000 µM, 17 | | |
| glutamate | $Cl^-$ | 0 µM, 7 | 3 µM, 8 | 30 µM, 8 | 100 µM, 8 | 1000 µM, 8 | 3000 µM, 8 | | | |
| | $Cl^-$ free | 0 µM, 7 | 3 µM, 7 | 30 µM, 8 | 100 µM, 8 | 1000 µM, 7 | 3000 µM, 4 | | | |
| | | $[Cl^-]$ mM, n | | | | | | | | |
| $[^3H]$taurine | $Na^+$ | 0.1 mM, 14 | 1 mM, 10 | 3 mM, 19 | 10 mM, 10 | 30 mM, 10 | 60 mM, 9 | 100 mM, 20 | 150 mM, 18 | |
| | $Na^+$ free | 0 mM, 10 | 1 mM, 10 | 10 mM, 20 | 30 mM, 10 | 60 mM, 9 | 100 mM, 20 | | | |
| L-$[^{14}C]$Aspartate | $Na^+$ | 0 mM, 10 | 1 mM, 10 | 10 mM, 8 | 100 mM, 7 | | | | | |
| | $Na^+$ free | 0 mM, 8 | 1 mM, 10 | 10 mM, 18 | 60 mM, 8 | 100 mM, 17 | | | | |
